# Keratin 14-dependent disulfides regulate epidermal homeostasis and barrier function via 14-3-3σ and YAP1

**DOI:** 10.1101/824219

**Authors:** Yajuan Guo, Krystynne A. Leacock, Catherine Redmond, Vinod Jaskula-Ranga, Pierre A. Coulombe

## Abstract

The type I intermediate filament (IF) keratin 14 (K14) provides vital structural support in basal keratinocytes of epidermis. Recent studies evidenced a role for K14-dependent disulfide bonding in the organization and dynamics of keratin IFs in skin keratinocytes. Here we report that knock-in mice harboring a cysteine-to-alanine substitution at codon 373 (C373A) in *Krt14* exhibit alterations in disulfide-bonded K14 species and a barrier defect secondary to enhanced proliferation, faster transit time and altered differentiation in the epidermis. A proteomics screen identified 14-3-3 as major K14 interacting proteins. Follow-up studies showed that YAP1, a transcriptional effector of Hippo signaling regulated by 14-3-3sigma in skin keratinocytes, shows aberrant subcellular partitioning and function in differentiating *Krt14*C373A keratinocytes. Residue C373 in K14, which is conserved in several other type I IFs, is thus revealed as a novel regulator of keratin organization and YAP function in early differentiating keratinocytes, with an impact on cell mechanics, homeostasis and barrier function in the epidermis.

## Introduction

The epidermis covering our skin and body is truly remarkable. It maintains a vital and multidimensional barrier to the outside environment, and to water, while renewing itself with rapid kinetics even under normal physiological conditions (Kubo et al. 2012). The mechanisms through which new progenitor cells are produced at the base of this stratified epithelium, pace themselves through differentiation, and maintain tissue architecture and function in spite of a high rate of cell loss at the skin surface are only partially understood (Wells and Watt 2018).

Keratin intermediate filaments are major protein constituents in epithelial cells and are encoded by a large family of 54 conserved genes that are individually regulated in a tissue-and differentiation-specific fashion (Schweizer 2006). An outstanding question is the extent to which keratin, and other types of intermediate filaments (IFs), participate in basic processes such as cell differentiation and tissue homeostasis. The type I keratin 14 (K14) and type II K5 co-polymerize to form the prominent IF apparatus that occurs in the progenitor basal layer of epidermis and related complex epithelia (Nelson and Sun 1983; Fuchs 1995). Two main roles have so far been ascribed to K5-K14 IFs. First, to provide structural support and mechanical resilience to keratinocytes in the basal layer of epidermis and related epithelia (Coulombe 1991; Vassar et al. 1991; Fuchs and Coulombe 1992). Second, to regulate the distribution of melanin with an impact on skin pigmentation and tone (Uttam et al. 1996; Betz et al. 2006; Gu and Coulombe 2007). Dominantly-acting missense alleles in either *KRT5* or *KRT14* underlie the vast majority of cases of epidermolysis bullosa simplex (EBS), a rare genetic skin disorder in which trivial trauma results in skin blistering secondary to the lysis of fragile basal keratinocytes (Bonifas 1991; Coulombe et al. 1991; Fuchs and Coulombe 1992; Lane et al. 1992). Such mutant alleles may also affect skin pigmentation (Gu and Coulombe 2007), establishing the relevance of both roles of K5-K14 in both healthy and diseased skin.

Structural insight gained from solving the crystal structure of the interacting 2B regions of corresponding rod domain segments in human K5 and K14 highlighted the presence of a trans-dimer, homotypic disulfide bond involving cysteine (C) residue 367 (C367) in K14 (Coulombe and Lee 2012). The C367 codon is conserved in *KRT14* orthologs of higher metazoans and in many other type I keratin genes expressed in skin or oral epithelia (Strnad et al. 2011; Lee et al. 2012) (**Fig. 1a,b**). Conspicuously, this residue occurs within a four-residue interruption (stutter) of the long-range heptad repeat within coil 2 of the central alpha-helical rod domain in virtually all IF proteins (Lee et al. 2012) (**Fig. 1b**). K14 C367-dependent disulfides indeed form in human and mouse skin keratinocytes (Lee et al. 2012). Follow-up studies revealed that K14-dependent disulfide bonding plays a role in the assembly, organization and steady state dynamics of keratin IFs in live skin keratinocytes (Feng and Coulombe 2015a; Feng and Coulombe 2015b). Loss of the stutter cysteine alters K14’s ability to become part of the dense meshwork of keratin filaments that arises in the perinuclear space of early differentiating keratinocytes (Lee et al. 2012; Feng and Coulombe 2015a; Feng and Coulombe 2015b). Here we report on studies involving a new mouse model from which we provide evidence that K14-dependent disulfide bonding regulates entry into differentiation and thus the balance between proliferation and differentiation through regulated interactions with 14-3-3 adaptor proteins and YAP1, a terminal effector of Hippo signaling (Schlegelmilch et al. 2011; Silvis et al. 2011; Sambandam et al. 2015). We also discuss evidence that this role likely applies to K10 and other type I keratins expressed in surface epithelia.

**Figure 1.**
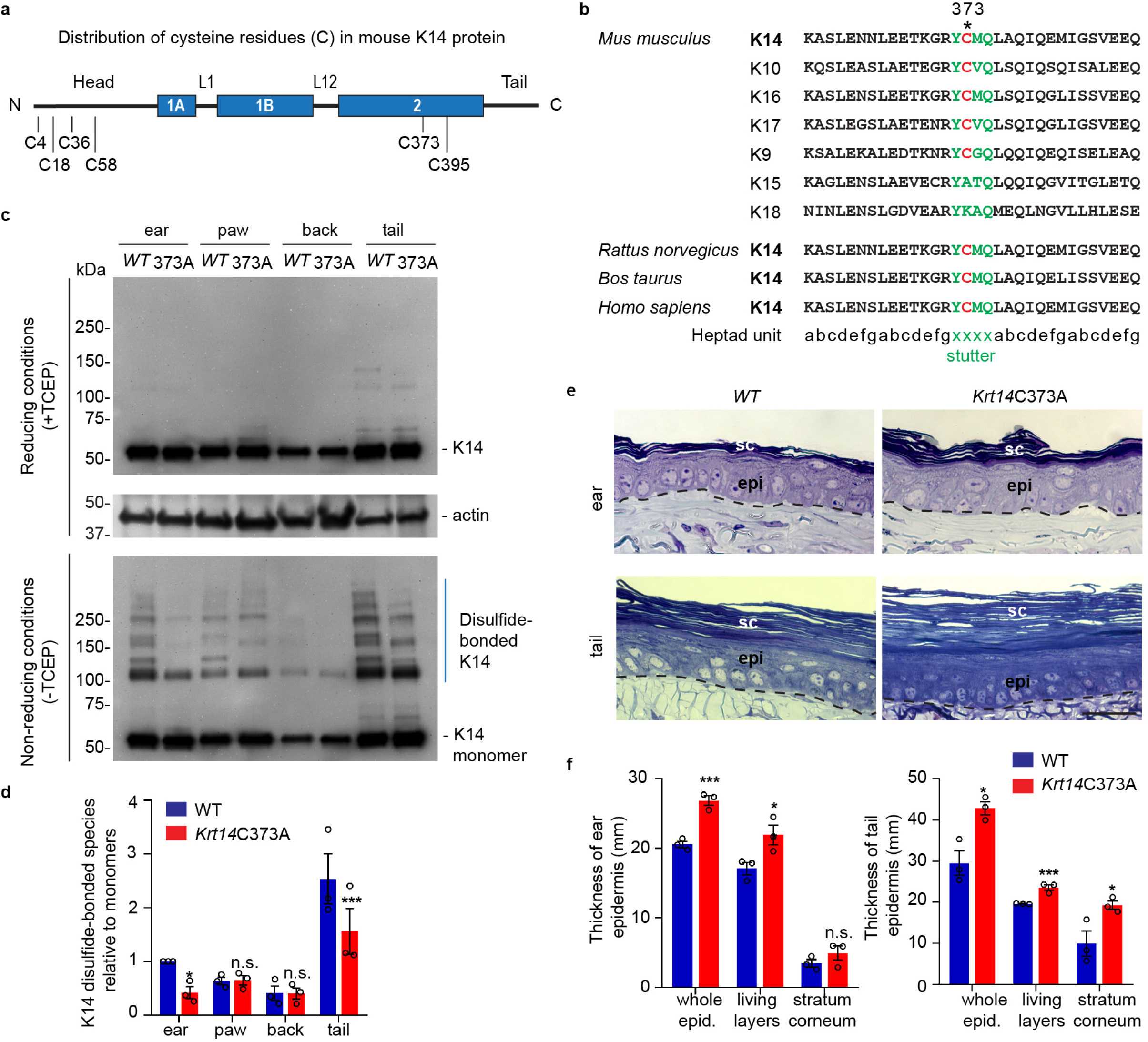
Decreased K14-dependent disulfide-bonded species and thickened epidermis in Krt14C373A mouse skin. a. Location of cysteine (C) residues in mouse K14 protein (C4, C18, C36, C58, C373, C395), in which N-terminal head and C-terminal tail domains are flanking the central □-helical rod domain (coils 1A, 1B and 2 (blue boxes) separated by linker s L1 and L12). b. Alignment of the sequence context flanking residue C373 in mouse K14 and in other mouse type I keratins (top) as well as in other species (bottom). The heptad units is shown at the bottom. “xxxx” makes the location of the stutter sequence (green letters). c. Immunoblotting analysis of total protein lysates from ear, paw, back skin, and tail skin from *WT* and *Krt14*C373A young adult mice subjected to SDS-PAGE electrophoresis under reducing (+TCEP) and non-reducing (-TCEP) conditions. d. Quantification of relative amounts of K14-dependent disulfides over monomers (see c). N=3 replicates. e. Toluidine blue-stained sections (1 mm thick from epoxy-embedded skin of young adult *WT* and *Krt14*C373A mice. f. Quantification of whole epidermal thickness (living epidermal layers and stratum corneum layers) in ear (left) and tail (right) skin of *WT* and *Krt14*C373A mice. 5 random fields were sampled for each of 3 mice per genotype. Data represent mean ± SEM. Student’s t test: n.s., no statistical difference; \**P* < 0.05; \*\*\**P* < 0.005. Scale bar, 20 μm.

## Results

To address the physiological significance of the stutter cysteine in K14, we generated *Krt14*C373A mutant mice using CRISPR-Cas9 technology (C373 in mouse corresponds to C367 in human K14; **Suppl. Fig. 1a,b**). *Krt14*C373A mice are born in the expected mendelian ratio, and are viable and fertile. While they are hardly distinguishable from *WT* littermates, *Krt14*C373A mice have a lower body weight when reaching adulthood (**Suppl. Fig. 1c**). Analysis of total skin proteins from several body sites showed that steady state levels of K14 protein are unaffected in *Krt14*C373A relative to *WT* skin. By contrast, the pattern of K14-dependent, high molecular weight disulfide-bonded species is markedly altered, given fewer species that occur at reduced levels (**Fig. 1c,d**). This is so especially in ear and tail skin (**Fig. 1c,d**), prompting us to focus on these two body sites in subsequent analyses. The residual K14-dependent disulfide bonding occurring in *Krt14* C373A mutant skin (**Fig. 1c,d**) likely reflects the participation of cysteines located in the N-terminal domain of K14 ((Feng and Coulombe 2015a); **Fig. 1a**). By histology, the epidermis of *Krt14*C373A mice is modestly but significantly thickened relative to *WT* in ear and tail skin (**Fig. 1e,f**; data not shown).

We next analyzed the skin barrier status in young adult *Krt14*C373A mice. Measurement of trans-epidermal water loss (TEWL) at the skin surface revealed an increase in *Krt14*C373A mice relative to *WT* control. This is so both at baseline (7.02±0.72 g/m^2^/h vs. 2.95±0.49 g/m^2^/h) and after topical acetone application (19.20±1.78 g/m^2^/h vs. 7.02±0.72 g/m^2^/h) (**Fig. 2a**), a standard challenge that puts the skin barrier under a mild and reversible stress (Denda et al. 1996). Skin barrier defects often trigger elevated expression of Danger-Associated Molecular Patterns (Lessard et al. 2013) (DAMPs, also known as alarmins). At baseline, DAMPs such as *S100A8, S100A9, MMP9*, and *Ptgs2* are upregulated by 4-fold or more at the mRNA level in *Krt14*C373A skin compared with *WT* (**Fig. 2b**), and these aberrant levels are even more robustly elevated after a topical acetone challenge to the barrier (**Fig. 2c**). Next we analyzed cornified envelopes (CEs) isolated from epidermis, given that they are key contributors to skin barrier function (Eckhart et al. 2013). CEs harvested from *WT* mice appear relatively uniform in size and shape, are mostly oval-shaped, and feature clear and smooth outlines (**Fig. 2d,e**). By contrast, CEs isolated from *Krt14*C373A mice are smaller (85% of the area and 87% of the circumference of *WT* CEs), jagged and less oval-shaped (aspect ratio of 1.3 compared to 1.1 in *WT*) (**Fig. 2d,e** and **Suppl. Fig. 1d,e**). Thus, the mild thickening and hyperkeratosis observed in epidermis are accompanied by significant defects in barrier function in *Krt14*C373A skin.

**Figure 2.**
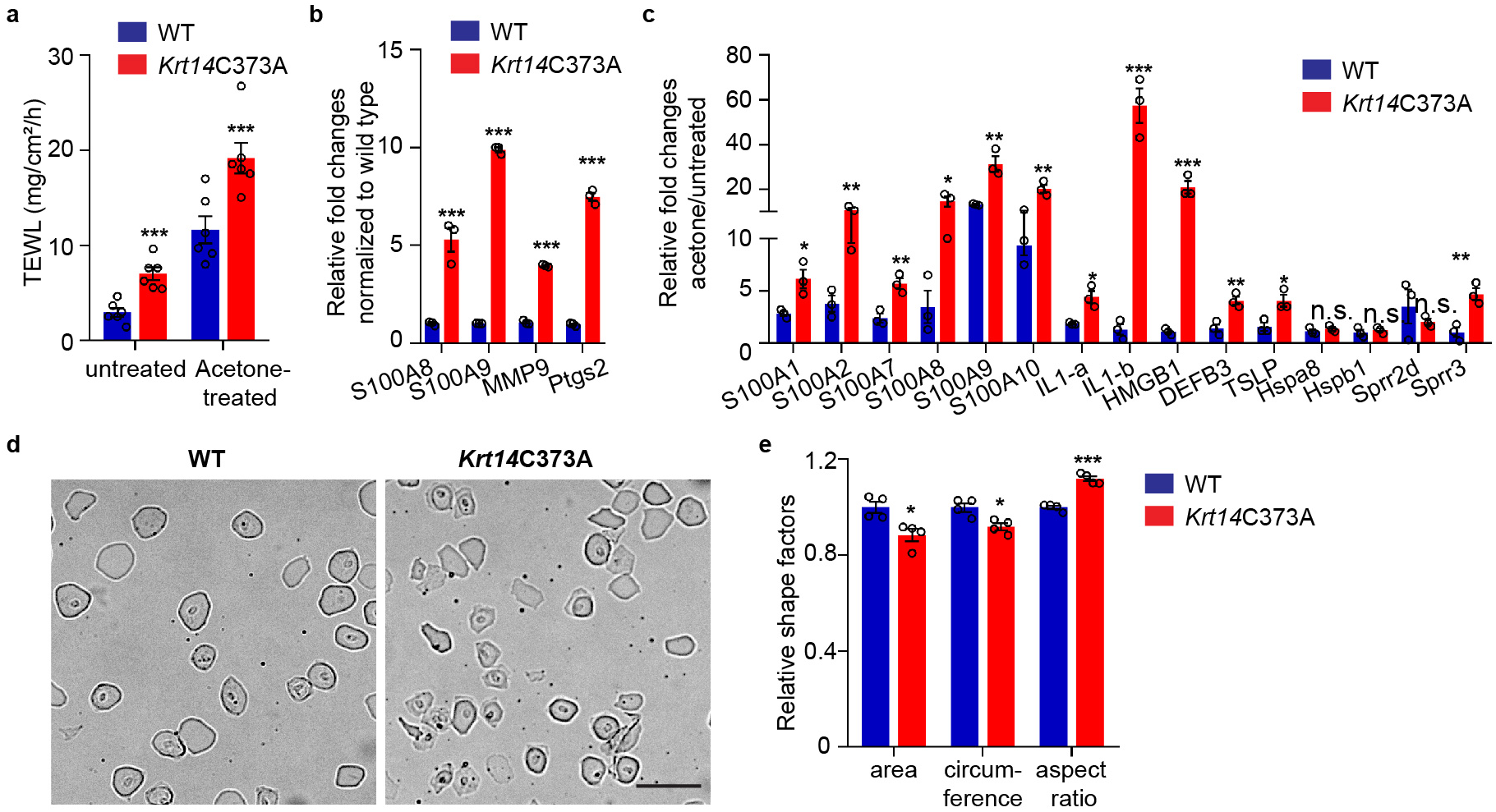
The epidermal barrier is defective in Krt14C373A skin. a. Trans-epidermal water loss measurements of *WT* and *Krt14*C373A ear skin at baseline and after acetone-induced barrier disruption. N=6 per sample. b. Relative fold change in mRNA levels (qRT-PCR) for Danger-Associated Molecular Patters (DAMPs) in *WT* and *Krt14*C373A skin at baseline. N=3 biological replicates. c. Relative fold change in mRNA levels (qRT-PCR) for DAMPs after acetone treatment. N=3 biological replicates. d. Representative images of cornified envelopes isolated from *WT* and *Krt14*C373A tail skin. e. Quantitation of surface area, circumference, and aspect ratio of isolated cornified envelopes in d. Approximately 100 CEs were counted for each of four mice. Data represent mean ± SEM. Student’s t test: \**P* < 0.05; \*\**P* < 0.01; \*\*\**P* < 0.005; n.s., no statistical difference. Scale bar, 100 μm.

We next assessed functional readouts related to proliferation, transit time, and apoptosis in *Krt14*C373A relative to *WT* epidermis. At 2 h after a single pulse of the nucleotide analog Edu (Chehrehasa et al. 2009), a significantly greater fraction of keratinocytes are labeled in the basal layer of *Krt14*C373A epidermis compared to *WT* (by ∼1.6 fold; p = 0.024)(**Fig. 3a,b**), indicating that proliferation is enhanced at baseline in mutant mice. The difference between genotypes is accentuated at 1 day after the pulse (>2-fold; p = 0.01). At that time, Edu-labeled nuclei occur in the suprabasal layers of epidermis in both genotypes (**Fig. 3a,b**), reflecting keratinocyte exit from the basal layer. Nuclear labeling remains high and stable in the basal layer of epidermis in both genotypes at the 3-day mark after the pulse, but a clear additional difference emerges in that the mutant epidermis exhibits a larger number of Edu-labeled nuclei in suprabasal layers. At the 7-day mark, the fraction of labeled cells in the basal layer has subsided in both genotypes but, again, the suprabasal epidermis of *Krt14*C373A skin shows far more labeled nuclei (**Fig. 3a,b**). This pulse-chase experiment establishes that, at baseline, keratinocytes in *Krt14*C373A epidermis show enhanced proliferation confined to the basal layer along with a faster pace of movement across the suprabasal layers as they progress through differentiation. A similar phenotype has been previously described for *Krt10* null mice (Reichelt and Magin 2002).

**Figure 3.**
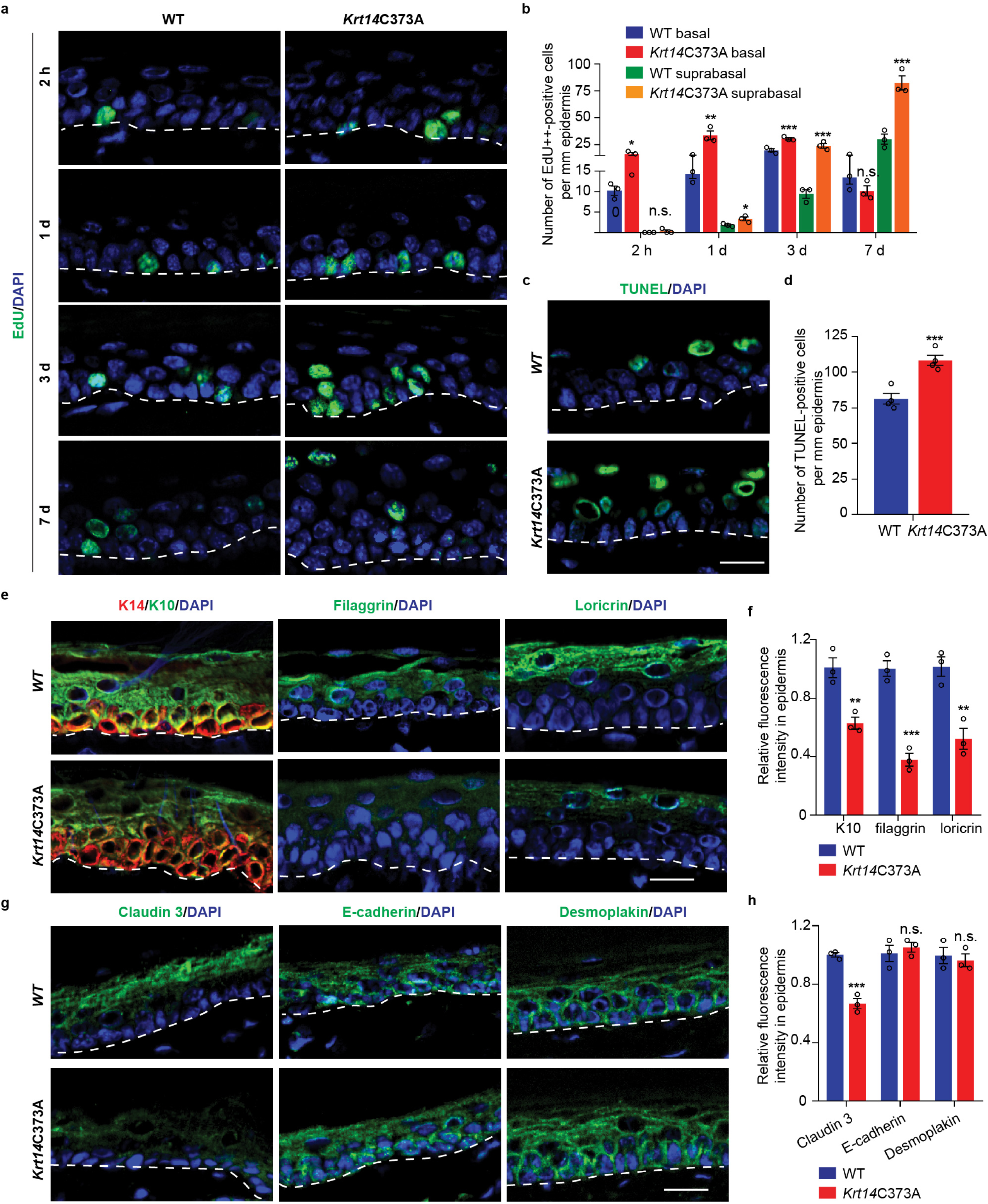
Altered tissue homeostasis and dysregulated keratinocyte differentiation in Krt14C373A skin. a. Indirect immunofluorescence for Edu in tail skin section from *WT* and *Krt14*C373A at 2 h, 1 d, 3 d, and 7 d after treatment with thymidine analog EdU. Nuclei as stained with DAPI (blue). Dashed lines depict the dermo-epidermal interface. b. Quantification of the number of EdU positive nuclei in basal and suprabasal layers per mm of epidermis. N=3 replicates for each sample. c. TUNEL staining in tail epidermis of young adult *WT* and *Krt14*C373A mice. d. Quantification of TUNEL-positive cells shown in frame c. N=4 mice per measurement. e. Indirect immunofluorescence for K14 (green), K10 (red), filaggrin, and loricrin from tail skin sections of *WT* and *Krt14*C373A mice. f. Quantification of relative fluorescence intensity of data shown in frame e, normalized to *WT*. N=3 mice per sample. g. Indirect immunofluorescence for claudin 3, E-cadherin and desmoplakin in tail skin sections from *WT* and *Krt14*C373A mice. h. Quantitation of relative fluorescence intensity in g. N=3 mice per sample. Data represent mean ± SEM. Student’s t test: \**P* < 0.05; \*\**P* < 0.01; \*\*\**P* < 0.005; n.s., no difference. Scale bars, 20 μm.

We also observed a greater frequency (∼3-fold) of TUNEL-positive nuclei in *Krt14*C373A epidermis compared to *WT*, with apoptotic cell death confined to the suprabasal compartment (**Fig. 3c,d**). When combined, the increases observed in the rate of keratinocyte proliferation in the basal layer and of the transit time and apoptosis in the suprabasal layers suggest that the modest increase observed in epidermal thickness (**Fig. 1e,f**) masks a more pronounced defect in epidermal homeostasis under baseline conditions in *Krt14*C373A mice.

We next sought confirmation that keratinocyte differentiation is altered in *Krt14*C373A skin. We examined the distribution of K14 (basal cell layer), K10 (early differentiation), and filaggrin and loricrin (late differentiation) using sections of tail skin from young adult mice. The signal for K10 was modestly decreased (∼37% reduction) while that for filaggrin and loricrin were markedly decreased (∼62% and 48% reduction, respectively) in *Krt14*C373A epidermis relative to *WT* (**Fig. 3e,f**). We also examined markers of tight junctions (claudin 3), adherens junction (E-cadherin), and desmosomes (desmoplakin) since differentiation is accompanied by dynamic and tightly coordinated rearrangements of intercellular junctions. Claudin 3, which is also related to differentiation, was decreased by ∼33%. The signals for E-cadherin and for desmoplakin appeared slightly increased (**Fig. 3g**) but this may reflect the modest epidermal thicknening in *Krt14*C373A epidermis (**Fig. 3h**). These observations suggest that the anomalies in epidermal homeostasis and skin barrier are accompanied by clear defects in terminal keratinocyte differentiation in the skin epidermis of *Krt14*C373A mice.

We previously showed that replacing Cys with Ala at position 367 in human K14 does not abrogate 10-nm filament formation but leads to a reduction in the perinuclear clustering of keratin filaments in cultured keratinocytes (Feng and Coulombe 2015b). Transmission electron microscopy of epoxy-embedded skin tissue sections was next used to assess whether similar changes occur *in vivo*. In basal keratinocytes of *WT* epidermis, keratin IFs typically occur as bundles near the nucleus. In *Krt14*C373A basal keratinocytes, however, keratin IFs are absent from the perinuclear region and appear redistributed towards the cell periphery (**Suppl. Fig. 2a,b**). There is no ultrastructural evidence of cell fragility in *Krt14*C373A epidermis (**Suppl. Fig. 2a,b** and data not shown). We also find that nuclei feature a more ellipsoid shape along with a greater frequency of cytoplasmic invaginations (by ∼1.4 fold in basal keratinocytes and by ∼1.7 fold in suprabasal keratinocytes, respectively) compared to *WT* controls; **Suppl. Fig. 2c**). These anomalies suggest that the defects in the perinuclear keratin IF network result in changes in the mechanical properties of the nucleus in *Krt14* C373A basal layer keratinocytes, thus extending previous live imaging observations of *Krt14* null mouse keratinocytes transfected with a C367A mutant in K14 (Feng and Coulombe 2015b).

To identify potential pathways regulated by K14-dependent disulfides, we performed K14 co-immunoprecipitation (co-IP) assays followed by mass spectrometry (MS) analysis in protein extracts prepared from newborn *WT* keratinocytes in primary culture in the presence of 1 mM Ca^2+^, a condition that induces keratinocyte differentiation (Hennings et al. 1980). This screen identified 14-3-3σ and other 14-4-3 isoforms as major interacting partners for K14 in *WT* cell cultures (**Fig. 4a** and **Suppl. Table 1)**. There is a strong precedent for interactions between 14-3-3 proteins and keratins, Including K18 (Liao and Omary 1996; Ku et al. 1998), K17 (Kim et al. 2006), which occur in a phosphorylation-dependent fashion. GO analysis of the top 100 MS-identified proteins (**Suppl. Table 1**) revealed, when excluding other keratin proteins, a marked enrichment for terms associated with organelle organization (p<8.7×10^−14^), rab protein signal transduction (p<1.0×10^−13^) and cellular protein localization (p<5.2×10^−13^). Independent co-immunoprecipitation assays confirmed that both endogenous 14-3-3σ and transfected HA tagged-14-3-3σ physically interact with both *WT* K14 and *Krt14*373A mutant protein in mouse keratinocytes in primary culture (**Fig. 4b** and data not shown). The yield in 14-3-3σ was reduced in the *Krt14*373A mutant samples (**Fig. 4b**), a finding that was confirmed by proximity ligation assays (see below).

**Figure 4.**
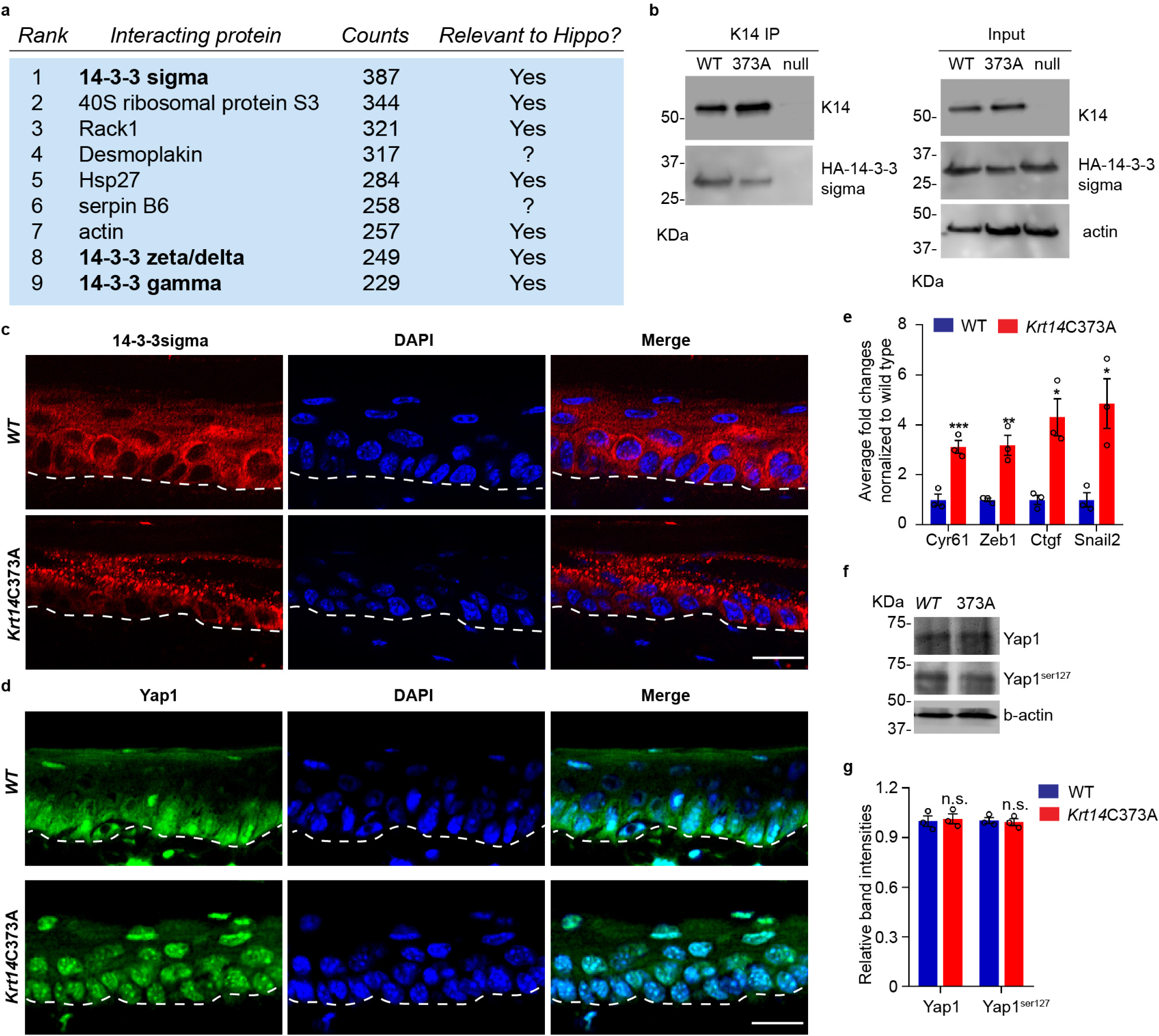
14-3-3σ interacts with K14, and abnormal localization 14-3-3σ and YAP in Krt14C373A epidermis. a. Top 9 most abundant non-keratin entries from a mass spectrometry screen for proteins interacting with K14 in wildtype newborn skin keratinocytes in primary culture (1 mM calcium, 4 days). Spectral counts and known relevance to Hippo signaling are indicated. See Suppl. Table 1 for full listing. b. Immunoprecipitation of K14 from *WT* or *Krt14*C373A skin keratinocytes in primary culture. Both K14 WT and, albeit to a lesser extent, the 373A mutant interact with HA-tagged 14-3-3σ. KDa, kilodalton. c. Indirect immunofluorescence for 14-3-3σ in *WT* and *Krt14* C373A tail skin sections. Dashed lines depict the dermo-epidermal interface. d. Indirect immunofluorescence for YAP in *WT* and *Krt14*C373A tail skin sections. e. Relative mRNA levels (qRT-PCR) for YAP target genes *Cyr61, Zeb1, Ctgf*, and *Snail2* in adult *WT* and *Krt14*C373A tail skin. N=3 biological replicates per genotype. f. Immunoblotting analysis for total YAP and YAP^Ser127^ in *WT* and *Krt14*C373A tail skin protein lysates. g. Quantification of relative protein levels shown in frame d. Data are mean ± SEM from 3 biological replicates. Student’s t test: \**P* < 0.05; \*\**P* < 0.01; \*\*\**P* < 0.005; n.s., no statistical difference. Scale bars, 20 μm.

14-3-3σ was deemed of interest because it regulates the proliferation and differentiation of keratinocytes in epidermis (Herron et al. 2005; Li et al. 2005). The latter is achieved in part by modulating the cellular localization of YAP (Li et al. 2005; Sun et al. 2015), a terminal effector of Hippo signaling (Schlegelmilch et al. 2011; Silvis et al. 2011; Sambandam et al. 2015). We next assessed the distribution of 14-3-3σ and YAP using indirect immunofluorescence of tissue sections prepared from *WT* and *Krt14*C373A tail skin. 14-3-3σ occurs mostly as aggregates in suprabasal keratinocytes of *Krt14*C373A epidermis, which is in striking contrast to the diffuse distribution observed in *WT* controls (**Fig. 4c**). Consistent with previous reports (Schlegelmilch et al. 2011; Sambandam et al. 2015), a strong signal for YAP occurs in both the nucleus and cytoplasm in basal keratinocytes, and otherwise YAP occurs as a weaker and diffuse signal in the cytoplasm (but is not seen in the nucleus) of suprabasal keratinocytes in *WT* epidermis (**Fig. 4d**). The situation is dramatically different in *Krt14*C373A epidermis, since YAP localizes preferentially to nuclei in both basal and suprabasal keratinocytes in a consistent fashion (**Fig. 4d**). The latter suggests that YAP-dependent gene expression may be altered in mutant mouse skin. Indeed, RT-qPCR assays show that the steady state levels for several known YAP target gene mRNAs, including *Cyr61, Zeb1, Ctgf* and *Snail2*, are markedly elevated in *Krt14*C373A relative to *WT* (**Fig. 4e**). By western immunoblotting, levels of endogenous YAP1 and Ser^127-^ phosphorylated YAP1 appear indistinguishable between *WT* and *Krt14*C373A skin (**Fig. 4f,g**), though a conclusive assessment of the status of these markers necessitates better reagents. Altogether these findings pointed to the misregulation of 14-3-3σ and YAP as likely contributors to the epidermal phenotype exhibited by *Krt14*C373A skin.

Next we asked whether the misregulation of YAP subcellular partitioning also occurs in primary culture. Keratinocytes were isolated from *WT* and *Krt14*C373A newborn pups, cultured in the absence or presence of calcium (Hennings et al. 1980), and analyzed. In the absence of calcium, keratinocytes preferentially proliferate and YAP is concentrated in the nucleus in both *WT* and *Krt14*C373A keratinocytes (**Fig. 5a,b**). After addition of calcium (1 mM), which triggers differentiation and mimics a suprabasal state (Hennings et al. 1980), 73% of *WT* keratinocytes lose their nuclear YAP signal whereas 95% of *Krt14*C373A keratinocytes exhibit predominantly nuclear YAP (**Fig. 5a,b**). Also, PLA assays yielded evidence for decreased interactions between 14-3-3σ and YAP in *Krt14*C373A keratinocytes in primary culture, relative to *WT* controls (**Fig. 5c,d**), thereby confirming the immunoprecipitation findings. Therefore, the abnormal retention of YAP to the nucleus in *Krt14*C373A keratinocytes is preserved outside of the skin tissue setting, suggesting that this trait is inherent to keratinocytes and manifested early during differentiation.

**Figure 5.**
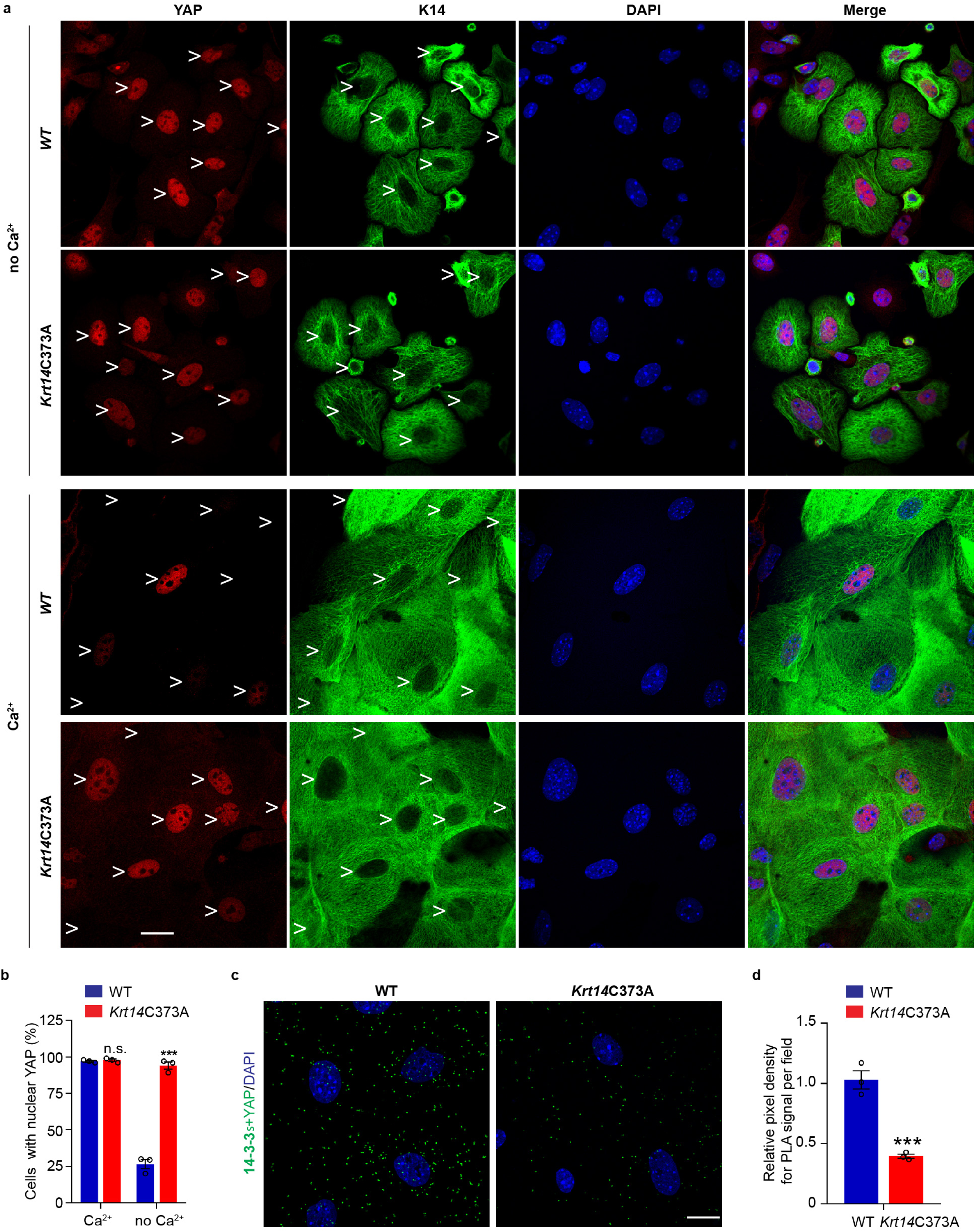
Localization and interaction of 14-3-3σ and YAP activities in Krt14C373A keratinocytes. a. Indirect immunofluorescence microscopy for YAP (red chromophore) and K14 (green chromophore) in *WT* and *Krt14*C373A newborn skin keratinocytes in primary culture in the absence and presence of 1 mM calcium (for 4 d). Arrowheads depict location of nuclei. b. Quantification of cells with nuclear YAP in frame a. N=3 biological replicates. Approximately 100 cells were counted for each genotype for each condition. c. Proximity ligation assay (PLA) for 14-3-3σ and YAP in *WT* and *Krt14*C373A newborn skin keratinocytes in primary culture in the presence of 1 mM calcium for 4 d. d. Quantification of relative fluorescence intensity in frame c. N=3 biological replicates. Data represent mean ± SEM. Student’s t test: n.s., no statistical difference; \*\*\**P* < 0.005. Scale bars, 20 μm.

The five cysteine residues present in human K14 are conserved in the mouse ortholog (Lee et al. 2012), and cysteines at positions 4, 40, and 367 in human K14 participate in disulfide bonding (Feng and Coulombe 2015a). We next asked whether the aberrant YAP nuclear localization observed in *Krt14*C373A mutant epidermis is Cys373-specific. Various GFP-K14 variants were transfected in *Krt14* null mouse keratinocytes in primary culture in the presence of 1 mM Ca^2+^, followed by YAP immunostaining via indirect immunofluorescence. Consistent with previous findings (Sambandam et al. 2015), only 19.5% of GFP-K14WT-expressing, calcium-treated keratinocytes exhibit nuclear YAP (**Suppl. Fig. 3a,b**). By contrast, cells expressing either a GFP-K14 cysteine free (CF) mutant or a GFP-C367A single mutant feature abnormally high nuclear retention of YAP (83.8% and 81.9%, respectively), compared to GFP-K14 expressing cells (**Suppl. Fig. 3a,b**). Restoring Cys367 in the K14-CF backbone (GFP-K14CF-C367) rescued the abnormal nuclear retention of YAP, since only 27.3% of transfected cells show YAP in the nucleus (**Suppl. Fig. 3a,b**). Given the established role of the stutter Cys in fostering the formation of a perinuclear network of keratin filaments (Lee et al. 2012; Feng and Coulombe 2015a; Feng and Coulombe 2015b), we infer from these new findings that this spatial organization is required for the proper regulation of YAP in keratinocytes.

In an effort to directly relate K14 to the activity of YAP1 as a transcriptional regulator, we next conducted luciferase reporter assays in a heterologous setting, HeLa cells in culture. Transfection of a *Cyr61* (*CCN1)* gene promoter-driven plasmid, previously shown to report on YAP transcriptional activity (Ma et al. 2017), led to a strong (∼6-fold) induction of luciferase activity over background activity in cells transfected with Renilla alone (**Suppl. Fig. 3c**). This activity was significantly attenuated by expression of WT K14 along with its natural partner WT K5 (**Suppl. Fig. 3c**). In striking contrast, luciferase activity was unaffected when K14 CF was co-expressed with WT K5 (**Suppl. Fig. 3c**). These findings provide support for a direct role for K14-containing filaments in negatively regulating YAP1, in a cysteine-dependent fashion.

YAP is a key effector of mechanosensing and mechanotransduction (Dupont et al. 2011; Benham-Pyle et al. 2015; Panciera et al. 2017), while K14 has been shown to provide vital mechanical support in basal keratinocytes of the epidermis (Coulombe 1991; Vassar et al. 1991; Fuchs and Coulombe 1992). Cells experiencing tension typically respond by forming more F-actin stress fibers heightened acto-myosin contraction, enhanced recruitment of α-catenin and vinculin to AJs (Leckband and de Rooij 2014; Yap et al. 2018), enhanced expression of lamin A/C (Swift et al. 2013), and increased frequency of binucleated cells (Cao et al. 2017). We next assessed the status of these mechanosensitive markers in skin tissue and in keratinocytes in primary culture. Relative to *WT* control, the signals for α-catenin and lamin A are markedly increased in *Krt14*C373A keratinocytes in epidermis in situ and in keratinocytes in culture (**Fig. 6a,b,c**). This is paralleled by the finding of prominent F-actin fibers and an increased staining intensity for phosphorylated myosin light chain II (pMLC Ser19) in *Krt14*C373A cells in primary culture (**Fig. 6e**). We also observed a greater incidence of multi-nucleated keratinocytes in Ca^2+^-treated *Krt14*C373A cultures compared to *WT* control (14.3% versus 2.8%; **Fig. 6d**). Collectively these findings are consistent with YAP misregulation (**Fig. 5**) and suggest that *Krt14*C373A mutant keratinocytes are defective in mechano-sensing and/or mechano-transduction.

**Figure 6.**
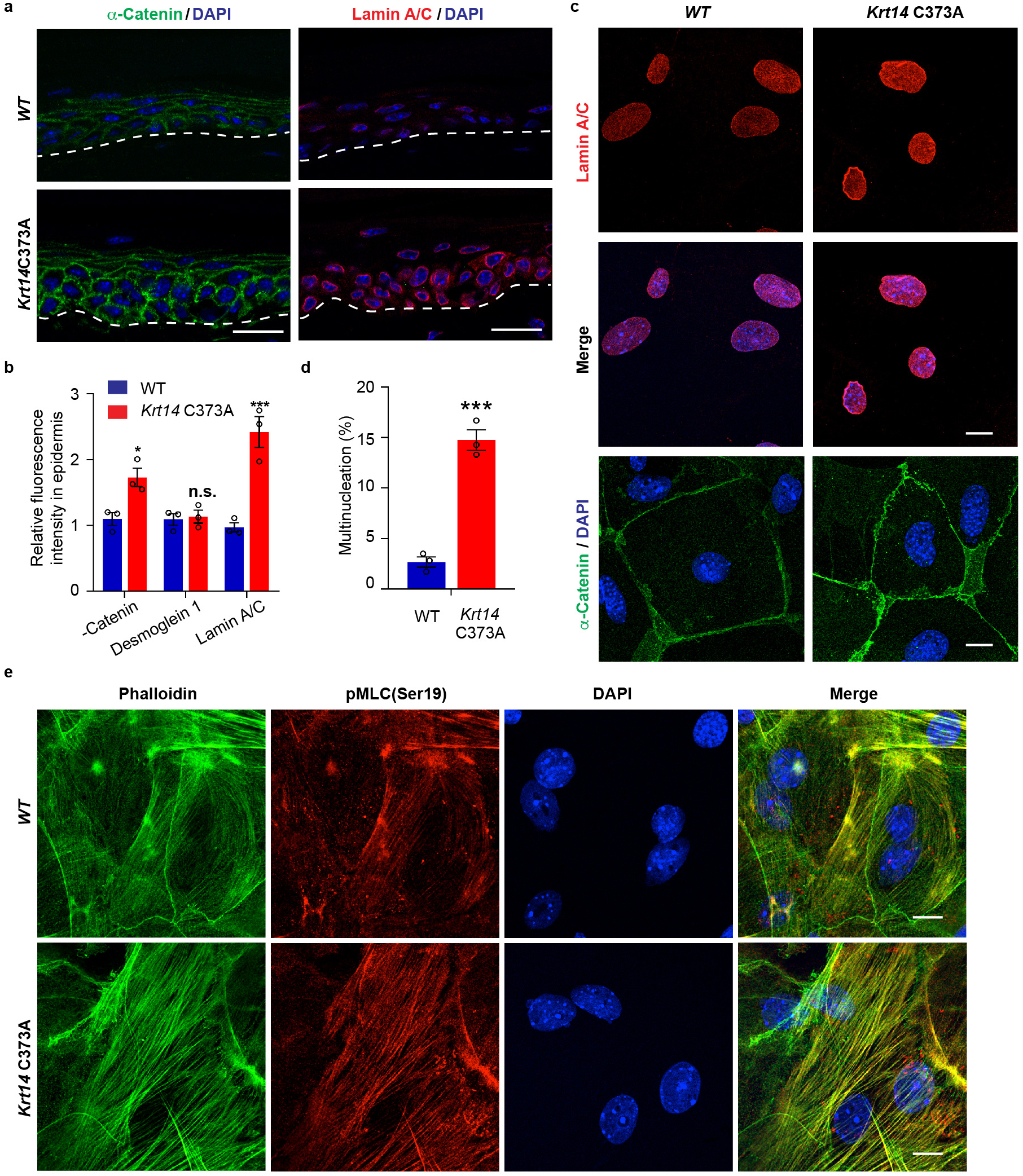
Altered mechanics in Krt14C373A skin and keratinocytes. a. Indirect immunofluorescence microscopy for *α*-catenin and lamin A/C in tail skin sections from young adult *WT* and *Krt14*C373A mice. Dashed lines depict the dermo-epidermal interface. b. Quantification of relative fluorescence intensity, as shown in frame a, in *Krt14*C373A relative to *WT* tissue. N=3 biological replicates. c. Indirect immunofluorescence for lamin A/C and *α*-catenin *WT* and *Krt14*C373A mutant newborn skin keratinocytes in primary culture (1 mM calcium for 4 days). d. Percentage of cells with multinucleation in *WT* and *Krt14*C373A keratinocytes cultured as described for frame c. N=3 biological replicates (total of 100 cells counted each time per genotype). e. Indirect immunofluorescence for (d) lamin A/C (e) and F-actin (green) and pMLC (Ser19) (red) in *WT* and *Krt14* C373A keratinocytes cultured ad described in frame c. Nuclei are stained with DAPI in frames a, c and e. Scale bars, 20 μm. Data in b and d represent mean ± SEM. Student’s t test: \**P* < 0.05; \*\*\**P* < 0.005; n.s., no statistical difference.

## Discussion

Our study establishes that residue cysteine 373 in mouse K14, which is conserved in the human ortholog and other mammals, regulates the organization of keratin IFs and the balance between keratinocyte proliferation and differentiation in epidermis *in vivo*, with an associated impact on skin barrier function. At a cellular and biochemical levels, loss of this cysteine residue results in profound alterations in: i) the pattern of K14-dependent disulfide bonding in epidermis; ii) the regulation of 14-3-3sigma and YAP in early-stage differentiating keratinocytes; and iii) mechanosensitive readouts in the epidermis in situ and keratinocytes in primary culture. When viewed in light of the literature (Miroshnikova et al. 2018; Nekrasova et al. 2018; Rubsam et al. 2018), our findings support the conclusion that site-specific K14-dependent disulfide bonding impacts cell architecture, mechano-sensing and Hippo signaling at an early stage of epithelial differentiation in the epidermis, possibly through the coordination of mechanical and biochemical cues as keratinocytes delaminate from the basal layer to enter the suprabasal compartment.

A model that conveys the significance our findings is given in **Fig. 7**. Consistent with the literature, the model posits that Hippo signaling is inactive in most keratinocytes in the basal layer of epidermis (Schlegelmilch et al. 2011; Beverdam et al. 2013; Sambandam et al. 2015; Totaro et al. 2017), YAP localizes to the nucleus of basal cells, reflecting a sub-threshold level of cellular crowding and/or integrin-mediated adhesion to the extracellular matrix (Panciera et al. 2017; Elbediwy and Thompson 2018), whereas 14-3-3σ occurs at low levels (Dellambra et al. 2000; Reichelt and Magin 2002; Kim et al. 2006) and K5-K14 filaments are organized in loose bundles that run alongside the nucleus (Coulombe et al. 1989; Lee et al. 2012). The model proposes that reception of differentiation-promoting cues rapidly triggers K14-dependent disulfide bonding and creates binding sites for 14-3-3σ on K5-K14 filaments, which are now organized into network of orthogonally-oriented filaments surrounding the nucleus. These events are paralleled by a major reorganization of cell-cell and cell-matrix adhesion and of F-actin and microtubule organization (Sumigray and Lechler 2015; Muroyama and Lechler 2017; Miroshnikova et al. 2018; Rubsam et al. 2018; Wickstrom and Niessen 2018), resulting in a redistribution of intracellular tension and/or compressive forces (Miroshnikova et al. 2018; Rubsam et al. 2018). The resulting sequestration of YAP1 to the cytoplasm activates Hippo signaling (**Fig. 7**). These findings reported here show that many of the features of this model are markedly disrupted in the absence of mouse K14’s cysteine 373, in vivo and in vitro.

**Figure 7.**
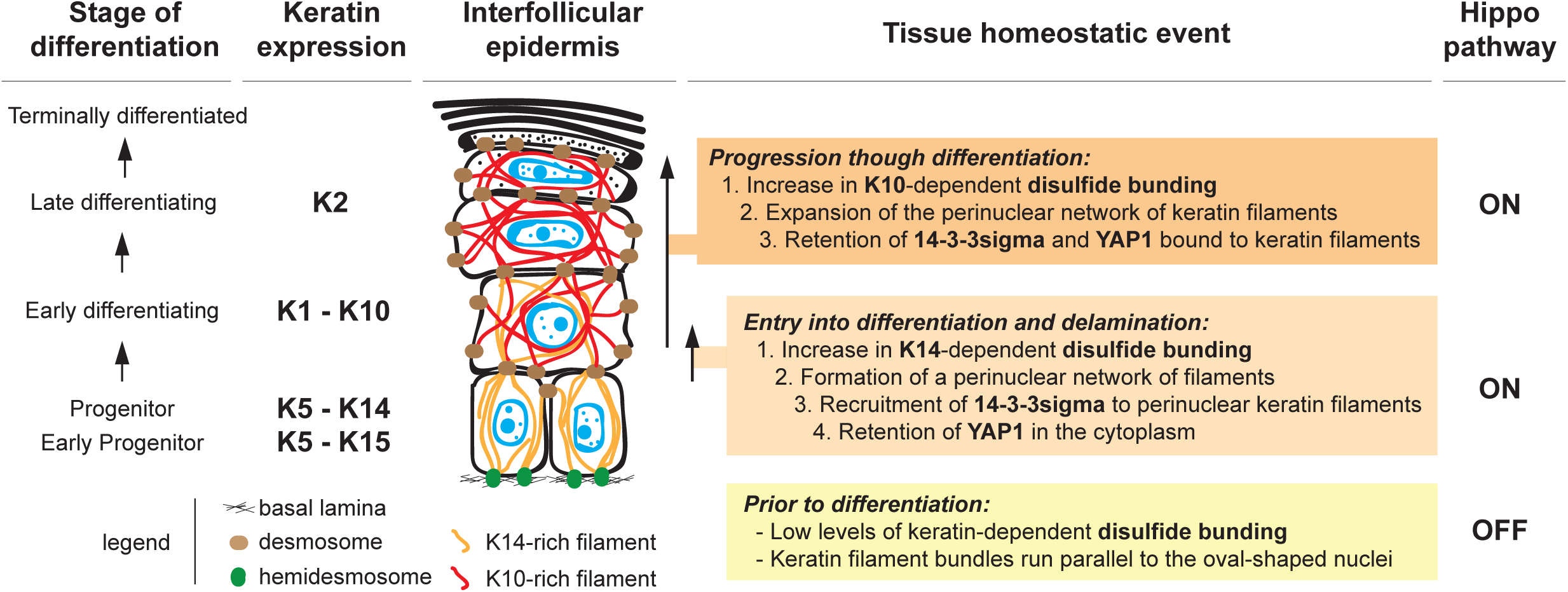
Model depicting the role of keratin-dependent disulfide bonding in integrating mechanical and signaling cues as keratinocytes initiate terminal differentiation in epidermis. Left to right: the stage of epidermal differentiation, keratin expression, epidermal morphology, and state of keratin filament organization are related to 14-3-3 binding, YAP1 subcellular partitioning, and Hippo activity status. The model proposes that initiation of terminal differentiation in late stage progenitor keratinocytes in the basal layer entails 1) the formation of K14-dependent disulfides via the conserved stutter cysteine in coil 2 domain; 2) reorganization of keratin filaments around the nucleus; 3) recruitment of 14-3-3 onto keratin filaments; and 4) effective sequestration of YAP1 in the cytoplasm, resulting in activation of Hippo signaling. The model proposes an identical role for the conserved cysteine in coil 2 of keratin 10, whisch is expressed early during terminal differentiation, thereby maintaining YAP1’s sequestration to the cytoplasm and active Hippo signaling. These changes are coupled to a redistribution of tension-related forces and cell-cell adhesion complexes as basal keratinocytes delaminate and move from the basal to the suprabasal compartment of epidermis (Miroshnikova et al. 2018; Nekrasova et al. 2018; Wickstrom and Niessen 2018). See text for additional details.

The model significantly extends reports linking 14-3-3σ to the regulation of YAP during terminal differentiation in epidermis (Sambandam et al. 2015; Sun et al. 2015). It suggests that the perinuclear keratin network may act as key docking site for 14-3-3/YAP complexes during keratinocyte delamination and differentiation, taking on a role that is held by integrin-based adhesion sites (Elbediwy et al. 2016) and adherens junctions (via alpha-catenin; (Schlegelmilch et al. 2011; Silvis et al. 2011) in the basal compartment of epidermis. Possibly, the perinuclear network of keratin filaments that arises in a keratin-dependent disulfide fashion could also afford protection to the nucleus and the genome during delamination (Lee et al. 2012), thus extending the established role of K5-K14 IFs in providing vital mechanical support in basal keratinocytes (Coulombe 1991; Ramms et al. 2013).

Many type I keratins expressed in epidermis feature a cysteine residue at the position corresponding to codon 373 in mouse K14 (Lee et al. 2012). Keratin 10, in particular, is a strong candidate for K14-like regulation of YAP1 subcellular partitioning and Hippo signaling in the epidermis. Three lines of evidence support this contention. First, Bunick and Milstone (Bunick and Milstone 2017) succeeded in crystallizing the interacting 2B segments of K1-K10, the keratin pairing characteristic of differentiating keratinocytes in epidermis (Fuchs 1995). The K1(2B)-K10(2B) structure (PDB 4ZRY) show an identical overall fold to our the original K5(2B)-K14(2B) structure (Lee et al. 2012), including the presence of a trans-dimer, homotypic disulfide bond mediated by the stutter cysteine (C401) in K10 (Bunick and Milstone 2017). Second, we already reported that K10 partakes in the formation of disulfide-dependent, dimer-sized species in skin keratinocytes (Lee et al. 2012), while others reported that K10 binds 14-3-3sigma (Wilker et al. 2007; Huang et al. 2010). Third, the *Krt10* null mouse exhibits an intriguing phenotype of hyperproliferation, faster keratinocyte transit time, and impaired differentiation in the epidermis which, molecularly, correlates with a marked upregulation of 14-3-3sigma and c-Myc (Reichelt and Magin 2002; Reichelt et al. 2004). *MYC* has since then been shown to be a *bona fide* YAP1 target gene (Schutte et al. 2014; Kim et al. 2017; Cai et al. 2018). Possibly, therefore, K10 could extend the role of K14 as keratinocytes progress through terminal differentiation (**Fig. 7**), and the disruption of this function through genetic mutations may play a role in the epidermis hyperproliferation that arises in epidermolytic hyperkeratosis and related conditions (Cheng et al. 1992; Chipev et al. 1992; Rothnagel et al. 1992; Kanitakis et al. 1993).

Several mechanisms could account for the functional interplay between K14-dependent disulfide bonding, 14-3-3σ, and the regulation of YAP’s subcellular partitioning. First, perinuclear enrichment of keratin IFs, which is promoted by K14-dependent disulfide bonding, might simply increase the local concentration of binding sites for 14-3-3σ and YAP near the nucleus (mass action law). Second, the occurrence of K14-dependent disulfides may create an optimal binding interface (or regulation thereof) for 14-3-3σ and YAP in the cytoplasm proximal to the nucleus. Third, the nucleus is known to function as a mechanosensor (Cao et al. 2017), and local forces impacted by the perinuclear network of keratin filaments could alter the mechanical gating of YAP across nuclear pores (Elosegui-Artola et al. 2017). These three mechanisms, and others, could act in combination. How K14, 14-3-3σ, YAP, and possibly other crucial effectors bind each other, how these interactions are regulated, and what is their significance have now emerged as open issues of high significance.

## Supporting information

Guo et al. Supplemental Information

## Acknowledgments

The authors are grateful to members of the Coulombe laboratory for support, to Samuel Black for technical support, to Dr. Steve Weiss and Dr. Yatrik Shah for reagents and advice, and to Dr. Roger Reeves, Chip Hawkins and the Transgenic Core facility at the Johns Hopkins University School of Medicine for the production of krt14 C373A mice. This work was supported by grant AR042047 to P.A.C. from the National Institutes of Health.

## Author contributions

Y.G. and P.A.C. designed the study, interpreted the findings and wrote the manuscript. V.J-R. designed strategies to target the *Krt14 gene* for mutagenesis and screen for the desired recombination event. Y.G. and P.A.C. oversaw the production of *Krt14*C373A mice, Y.G., K.L. and C.R. performed experiments.

## Methods

### Animals

All mouse studies were reviewed and approved by the Institutional Animal Use and Care Committee (IACUC) at both Johns Hopkins University and the University of Michigan. *WT* and *Krt14*C373A mice (C57BL/6 strain background) were maintained under specific pathogen-free conditions and fed rodent chow and water ad libitum. Male and female C57Bl/6 mice of 2-3 months of age (young adults) were used for all studies unless indicated otherwise.

### Generation of *Krt14*C373A mice using CRISPR-Cas9 technology

*Krt14*C373A mice were generated using the RNA-guided CRISPR-Cas9 system as described (Wang et al. 2013). A guide RNA (gRNA) was selected and designed according to a gRNA CRISPR design tool (http://crispr.technology) (Jaskula-Ranga 2016). Briefly, a Cas9 target site (GGGCCAGCTGCATGCAGTAACGG; with the PAM motif underlined) was selected based on having a cut site proximal to codon C373 and low-predicted off-targets. Oligonucleotides were used to clone the target into pT7gRNA, and the plasmid was amplified and linearized prior to T7 transcription. The gRNA was transcribed in vitro and purified prior to injection. The homology directed repair (HDR) template was purchased as a 183-nt single stranded Ultramer (IDT), and encoded a TGC (Cys) to GCA (Ala) mutation at codon 373 of the mouse K14 coding sequence. The gRNA, Cas9 mRNA, and HDR template were co-injected into C57Bl/6 zygotes by the JHU Transgenic Core Facility. Potential transgenic founders were screened using restriction digestion of PCR product extending beyond the repair template oligonucleotide and findings were confirmed by direct DNA sequencing (data not shown). Several founders exhibited the desired recombination event, either as homozygotes or heterozygotes. Two male homozygotes founders were selected and independently backcrossed through matings to C57Bl/6 *wildtype* females for two generations to eliminate potential off-target effects. The *Krt14*C373A homozygotes used in this study were from *Krt14*C373A het × het breedings (for body weight measurements and epidermal thickness measurements) or hom × hom breedings (other experiments). The two lines analyzed exhibited consistent and identical phenotypes.

### Topical acetone treatments of *Krt14*C373A mice

The left ears of age-matched *WT* and *Krt14*C373A mice (2-3 months old) were topically treated with 40 μl acetone twice daily for 7 days (Denda et al. 1996). The volume of acetone applied was split equally between the dorsal and ventral sides of the ear. The right ear (same mice) was left untreated as control. Mice were anesthetized during acetone treatment as per IACUC standards. Immediately after the last treatment, mice were euthanized and tissue harvested for analysis.

### Transepidermal water loss (TEWL) measurements

Mice were anesthetized using isoflurane (delivered by inhalation) during TEWL measurements. Readings were obtained using a TEWAMETER TM300 (Courage and Khazaka, Köln, Germany) from adult *WT* and *Krt14*C373A mice at baseline and after the last topical treatment with acetone. Measurements were made from the dorsal side of ear skin. The TM300 probe was warmed for 2 minutes prior to each measurement, and held on the area of interest for a minimum of 30 reads until the alpha level was below 0.2, per the manufacturer’s instructions.

### Measurement of cell proliferation through EdU labelling

EdU (A10044, Thermo Fisher Scientific) was prepared in PBS buffer at 10 mg/ml PBS and injected intraperitoneally into mice at a dose of 50 mg/kg body weight. Tail skin was harvested from anesthetized mice at 2 h, 1 d, 3 d, and 7 d after injection and processed for immunofluorescence staining. EdU staining was performed using the Click-iT Plus EdU Alexa Fluor 488 Imaging Kit (catalog no. C10637, Thermo Fisher Scientific).

### Immunofluorescence staining of skin tissue sections

For indirect immunofluorescence staining, ear or tail samples were surgically harvested and immediately submerged into optimal cutting temperature (O.C.T.) media (25608-930, VWR Scientific), flash frozen on dry ice, and stored at –40°C until sectioning. 5 μm cryosections were cut in a specific and consistent tissue orientation in all experiments. Cryosections were allowed to thaw in PBS buffer at room temperature and incubated with primary antibodies followed by Alexa Fluor–conjugated secondary antibodies (Thermo Fisher Scientific), counterstained in DAPI (1;5, 000 in PBS; D1306, Thermo Fisher Scientific), and mounted in FluorSave Reagent mounting medium (345789, Calbiochem) for indirect immuno-fluorescence (Hobbs et al. 2015; Kerns et al. 2016). Imaging was performed using either a a Zeiss fluorescence microscope with an Apotome attachment or a Zeiss LSM 800 confocal microscope. All experimental and control preparations were imaged under identical exposure conditions, and quantified using the ImageJ software (NIH). The primary antibodies used are listed in Suppl. Table 3. TUNEL staining for apoptotic cells was performed using the TUNEL enzyme (11767305001, Roche applied Science) and TUNEL label mix (11767291910, Roche applied Science) as recommended by the manufacturer.

### Isolation and analysis of cornified envelopes (CEs)

CEs were isolated from dorsal ear and tail tissue from age-matched male *WT* and *Krt14*C373A mice. To separate dorsal from ventral ear tissue, we followed the Murine Skin Tissue Transplant protocol (Garrod and Cahalan 2008). Extraction and preparation of CEs were performed using a protocol described by Kumar et al. (Kumar et al. 2015). Briefly, adult mouse ear skin or adult mouse tail skin (1cm length) were boiled at 95° C (in place of hot water bath) for 20 min in 2 ml CE isolation buffer containing 20 mM Tris-HCl (pH 7.5), 5 mM EDTA, 10 mM dithiothreitol (DTT), and 2% sodium dodecyl sulfate (SDS). Half of the extracted sample (1ml) was flash frozen and stored for future studies. CEs were extracted from the remaining (1ml) portion of the CE isolate. Samples were centrifuged for 5-minutes at 5, 000 × g, rinsed in CE isolation buffer with 0.2% SDS, re-pelleted, resuspended in 250 μl of washing buffer, and stored at 4°C until seeded. For morphological evaluation, CE isolates from dorsal ear and tail skin were seeded on glass slides at a concentration of 1.5 × 10^6^ CEs and 6 × 10^6^ CEs, respectively, covered with a thin cover glass, and then imaged. CEs were isolated from 4 mice per genotype. Analysis of the area, circumstance, and aspect ratio (longest axis to the shortest axis) of CEs was performed using ImageJ software.

### Transmission electron microscopy

Ear tissue from 2-3-month old *WT* and *Krt14*C373A littermates was surgically harvested, minced, and fixed overnight at 4 °C in 2% formaldehyde/2% glutaraldehyde in 0.1 M cacodylate buffer at pH 7.4. Samples were post-fixed in osmium tetroxide, counter-stained with uranyl acetate, and embedded in epoxy resin as previously described(Lessard et al. 2013). Thin sections were cut (50-70 nm thick), counter-stained with uranyl acetate and lead citrate, and examined using a Hitachi HU-12A transmission electron microscope. Toluidine blue-stained thick sections (1 μm thick) were used for morphological analyses at the light microscope level.

### RNA harvest, cDNA synthesis, and quantitative RT-PCR

RNA was harvested using TRIzol reagent (15596018, Thermo Fisher Scientific) and purified using the Nucleospin RNA kit (740955.250, Machery Nagel) according to the manufacturers’ instructions. Concentration and purity for RNA samples were assessed by spectrophotometry. 1.0 μg RNA was reverse-transcribed with the iScript cDNA Synthesis kit (1708891BUN, Bio-Rad Laboratories) using the manufacturer’s protocol. qRT-PCR was performed using iTaq Universal SYBR master mix (1725121, Bio-Rad Laboratories) on the CFX96 qRT-PCR apparatus (Bio-Rad Laboratories) as described (Hobbs et al. 2015; Kerns et al. 2016). The following program was used for all qRT-PCR reactions: denaturation step at 95°C for 5 min, 40 cycles of PCR (denaturation at 95°C for 10 s, annealing and elongation at 55°C for 30 s). No template or no reverse transcriptase controls, standard curves and a melt curve were included on every PCR plate. Normalized expression values from qRT-PCR data were calculated using Microsoft Excel by first averaging the relative expression for each target gene (2^−(Cq target gene – Cq reference gene)^) across all biological replicates and then dividing the relative expression value for the experimental condition by that for the control condition (2^−(ΔCq experimental – ΔCq control)^). Error bars were derived from the standard error of the mean (SEM) of the normalized expression values across all biological replicates. Normalized expression values for each target gene in all qRT-PCR experiments were derived from at least three biological replicates. Relative quantifications or fold changes of target mRNAs were calculated after normalization of cycle thresholds with respect to the reference gene β-actin. A list of all oligonucleotide primers used for custom qRT-PCR is provided in Suppl. Table 4.

### Isolation of skin keratinocytes for primary culture, calcium-induced differentiation and immunofluorescence stainings

Keratinocytes from 1 or 2-day old C57Bl/6 newborn mouse skin were isolated as described (Wang et al. 2016), and cultured in FAD medium (low calcium, 0.07mM) for 1 day. Calcium switch experiments (Wang et al. 2016) were performed by switching to FAD medium supplemented with with 1 mM CaCl_2_. Keratinocytes were harvested for analysis at 4 days or at 36 h after calcium switch as indicated in figure legends. For immunofluorescence, keratinocytes were fixed in 4% paraformaldehyde (PFA), blocked in 10% normal goat serum/0.1% Triton X-100/PBS for 1hr at room temperature, incubated in primary antibody solution for 1hr, washed in PBS, incubated in Alexa Fluor–conjugated secondary antibodies (Thermo Fisher Scientific), counterstained in DAPI (D1306, Thermo Fisher Scientific), and mounted in FluorSave Reagent mounting medium (345789, Calbiochem). Proximity ligation assay was performed according to the manufacturer’s protocol (Duolink in Situ PLA, Sigma-Aldrich). F-actin was stained using the Alexa Fluor 488 Phalloidin (A123791, Thermo Fisher Scientific) according to the manufacturer’s protocol. Micrographs were acquired using the Zeiss LSM 800 confocal microscope (Carl Zeiss Microscopy). Representative images from at least 3 independent experiments were shown. All images were and quantified by ImageJ software (NIH).

### Nucleofection of newborn mouse skin keratinocytes in primary culture

*Krt14*^-/-^ skin keratinocytes(Feng and Coulombe 2015a; Feng and Coulombe 2015b) were cultured in FAD medium. pBK-CMV His-GFP-K14WT or cysteine variants (Feng and Coulombe 2015a; Feng and Coulombe 2015b) were transfected into *Krt14*^-/-^ skin keratinocytes using P1 Primary Cell 4D-Nucleofector™ X Kit (V4XP-1024, Lonza). After nucleofection, cells were plated on collagen-coated coverglass and processed for analysis. For co-immunoprecipitation, HA-14-3-3σ (11946, Addgene) was transfected into skin keratinocytes in primary culture using the P1 Primary Cell 4D-Nucleofector™ X Kit (V4XP-1024, Lonza).

### Co-immunoprecipitation, protein gel electrophoresis, and mass spectrometry analysis

*WT* and *Krt14*C373A keratinocytes in primary culture were washed with PBS and lysed in cold Triton lysis buffer supplemented with Empigen (1% Triton X-100; 2% Empigen; 40 mm Hepes, pH 7.5; 120 mm sodium chloride; 50μ MN-ethylmaleimide; 1 mm EDTA; 1 mm phenylmethyl-sulfonyl fluoride; 10 mm sodium pyrophosphate; 1 μg/ml each of chymostatin, leupeptin, and pepstatin; 10 μg/ml each of aprotinin and benzamidine; 2 μg/ml antipain; 1 mm sodium orthovanadate; and 50 mm sodium fluoride). Protein concentration was determined using the Bio-Rad protein assay (Bio-Rad Laboratories) with bovine serum albumin (Thermo Fisher Scientific) as a standard. For immunoprecipitation, aliquots of cell lysate were incubated with a K14 antibody, and immune complexes were captured using the Protein G Sepharose (17-0618-01, GE Healthcare). Samples for gel electrophoresis were prepared in Laemmli Sample Buffer (LDS) sample buffer (1610747, Bio-Rad) in the presence of 20 mM tris(2-carboxyethyl)-phosphine (TCEP) (77720, Thermo Fisher Scientific) and incubated at room temperature for 1hr to reduce disulfide bonds. Non-reducing lysates were prepared directed in LDS sample buffer. Equal amounts of IP samples were resolved by 4–15% precast polyacrylamide gels (456-1084, Bio-Rad) and stained using a Silver Stain Kit (24612, Thermo Fisher Scientific). Bands of interest, along with a control area, were excised and analyzed by routine tandem mass spectrometry at the Johns Hopkins Mass Spectrometry Core. Mass spectrometry data were searched with Mascot 2.6.1 (Matrix Science) via Proteome Discoverer 2.2 (Thermo) against the RefSeq2017_83_mus_musculus Proteins database. Proteins with a false discovery rate (FDR) lower than 1% and with at least 2 identified peptides were reported as positive.

### Preparation of cell lysates, protein gel electrophoresis, and immunoblotting analysis

Cells or minced tissue were lysed in cold urea lysis buffer (pH 7.0, 6.5M urea, 50mM Tris-HCl, 150mM sodium chloride, 5mM ethylenediaminetetraacetic acid (EDTA), 0.1% Triton X-100, 50μM N-ethylmaleimide, 1mM phenylmethanesulfonyl fluoride (PMSF), 1μg/mL each of cymostatin, leupeptin, and pepstatin, 10μg/mL each of aprotinin and benzamidine, 2μg/mL antipain, and 50mM sodium fluoride). Protein concentration of the lysates was determined using Bradford protein assay (Bio-Rad) with bovine serum albumin as a standard. Samples for gel electrophoresis were prepared in LDS sample buffer (1610747, Bio-Rad) in the presence of 20 mM TCEP and incubated at room temperature for 1 h to reduce disulfide bonds. Non-reduced lysates were prepared directed in LDS sample buffer. Equal amounts of cell lysates were resolved by 4–15% precast polyacrylamide gels (Bio-Rad) and transferred to nitrocellulose membrane (0.45μm, Bio-Rad), and immunoblotted with the indicated antibodies followed by HRP-conjugated goat anti–mouse IgG or anti–rabbit IgG or rabbit anti–chicken IgY (Sigma-Aldrich) and Super Signal West Pico Chemiluminescent Substrate (PI34080, Thermo Fisher Scientific) or Amersham ECL Select Western Blotting Detection Reagent (RPN2235, GE Healthcare). Signals were detected using the FluorChem Q imaging system (Protein Simple). The ImageJ software (NIH) was used for Western blot signal quantitation.

### Luciferase assays

Renilla luciferase control plasmid pRL-TK (Promega, E2241), YAP activity responsive Firefly luciferase plasmid Cyr61-luc (Ma et al. 2017), expression plasmid of human keratin 5 (K5), and expression plasmids of wildtype and cysteine-free (CF) keratin 14 (K14) (Feng and Coulombe 2015a) were transfected into HeLa (ATCC) cells using SE Cell Line 4D X Nucleofector™ Kit S (V4XC-1032) with setting DS-138. After Nucleofection, cells were plated across six wells of a black matrix 96-well plate for each parameter. HeLa cells were transfected such that the cell density in each well was 30-40% the following morning. Firefly and Renilla luciferase activities were measured using Promega Dual Luciferase Reporter Assay System (Promega, PR-E1910). Firefly relative light unit (RLU) was normalized to internal Renilla RLU per well. Three biological replicates of normalized Firefly RLUs were pooled, and the means of each parameter were compared using a Mann-Whitney test. Data displayed was transformed by dividing individual RLUs of each parameter by the mean of pRL-TK alone and subjected to statistical analysis.

### Computational prediction of protein motifs

The predicted mouse K14 protein sequence (UniProtKB Q61781) was analyzed using publicly accessible algorithms written to predict 14-3-3 binding sites and phosphorylation events, including 14-3-3-Pred (Madeira et al. 2015) and Scansite 4.0 (Obenauer et al. 2003).

### Graphing and statistics

All graphs convey mean ± SEM values calculated using the Microsoft Excel software 2016 (Microsoft Office) or Prism software version 7 (GraphPad Software, Inc.). For comparisons between datasets, the Student’s t test (tails = 2) was used, and statistically significant p-values are indicated in figures and figure legends.

