## Supplementary material for "Keratin 14-dependent disulfides regulate epidermal homeostasis and barrier function via 14-3-3σ and YAP1": Guo et al. Supplemental Information

**List of elements:**

**Supplementary Figure 1**
***Generation of Krt14C373A mice and skin barrier measurements.*** a. Schematic diagram of the strategy used to generate *Krt14*C373A mice using the Crispr/Cas9 system. sgRNA, single guide RNA; PAM, protospacer adjacent motif. b. Sanger sequencing showing the TGC to GCA transversion at codon 373 (cysteine to alanine) in the *Krt14* gene. c. As young adult *Krt14*C373A males show a modestly reduced body mass relative to *WT* littermates. N=7 for each genotype. d. Representative microscopic images of cornified envelopes isolated (CE) from *WT* and *Krt14*C373A ear skin. Bar, 100 µm. e. Quantitation of surface area, circumference, and aspect ratio of CEs. N=4. Approximately 100 CE pieces were measured per mouse. Data represent mean ± SEM. Student's t test: n.s., no statistical difference; *$P < 0.05$.

**Supplementary Figure 2**
***Ultrastructural changes and abnormal nuclei in Krt14C373A keratinocytes.*** a. Transmission electron micrograph of ear skin epidermis from *WT* and *Krt14*C373A mice. b. High magnification images of representative cells in frame a. Dotted lines mark the dermo-epidermal interface. Asterisks depict areas showing gaps between keratin filaments (kf) and the nucleus (Nu) in mutant *Krt14*C373A keratinocytes. Arrowheads depict cytoplasmic invaginations into the nucleus in *Krt14*C373A keratinocytes. Scale bar, 5 µm. c. Quantification of frequency of nuclear invaginations (n=14 nuclei from 2 biological replicates for each genotype). Data represent mean ± SEM. Student's t test: ***$P < 0.005$.

**Supplementary Figure 3**
***Localization of YAP is specifically regulated by cysteine residue 367 in human K14.*** Newborn skin keratinocytes from *Krt14* null mice were seeded in primary culture, transiently transfected with either GFP-K14 WT, GFP-K14CF, GFP-K14C367A or GFP-K14CF-C367 constructs, and cultured in 1 mM calcium for 36 h prior to analysis. a. Indirect immunofluorescence showing successfully transfected and viable cells (see asterisks). Scale bar, 20 µm. b. Quantitation of the percentage of GFP-K14 transfected cells with nuclear YAP staining. N=3 biological replicates. Approximately 100 cells were counted for each genotype each time. Data represent mean ± SEM. Student's t test: ***$P < 0.005$; n.s., no difference. a. outcome of Lucirase assatys in HeLa cells transfected with a *Cyr61*(*CCN1*)-Luciferase reported constructs. Data were normalized with regard to transfection efficience and signal obtained with pRL-TK vector control. Data represent mean ± SEM. Statistical analysis was performed on non-transformed data: **$P< 0.01$, *$P<0.05$, n.s., non-significant.

**Supplementary Table 1**
***Mass spectrometric screen for proteins interacting with wildtype K14 in WT newborn skin keratinocyte in primary culture.*** List of all proteins with at least 35 spectral counts (cumulative over three biological replicates). Gene symbols are provided at left. WT1, WT2, WT3, WT4 and WT5 represent different regions of silver stained protein electrophoretic gels subjected to MS analysis (data not shown). Keratin protein entries were expected to dominate this screen, and are listed using blue lettering. 14-3-3 protein isoforms are listed using red lettering. The top non-keratin protein from this screen is 14-3-3sigma. Relate to Figure

**Supplementary Table 2**
***List of antibodies used in this study.***

**Supplementary Table 3**
***List of oligonucleotide primers used in this study.***

### Guo et al. Supplementary Figure 1

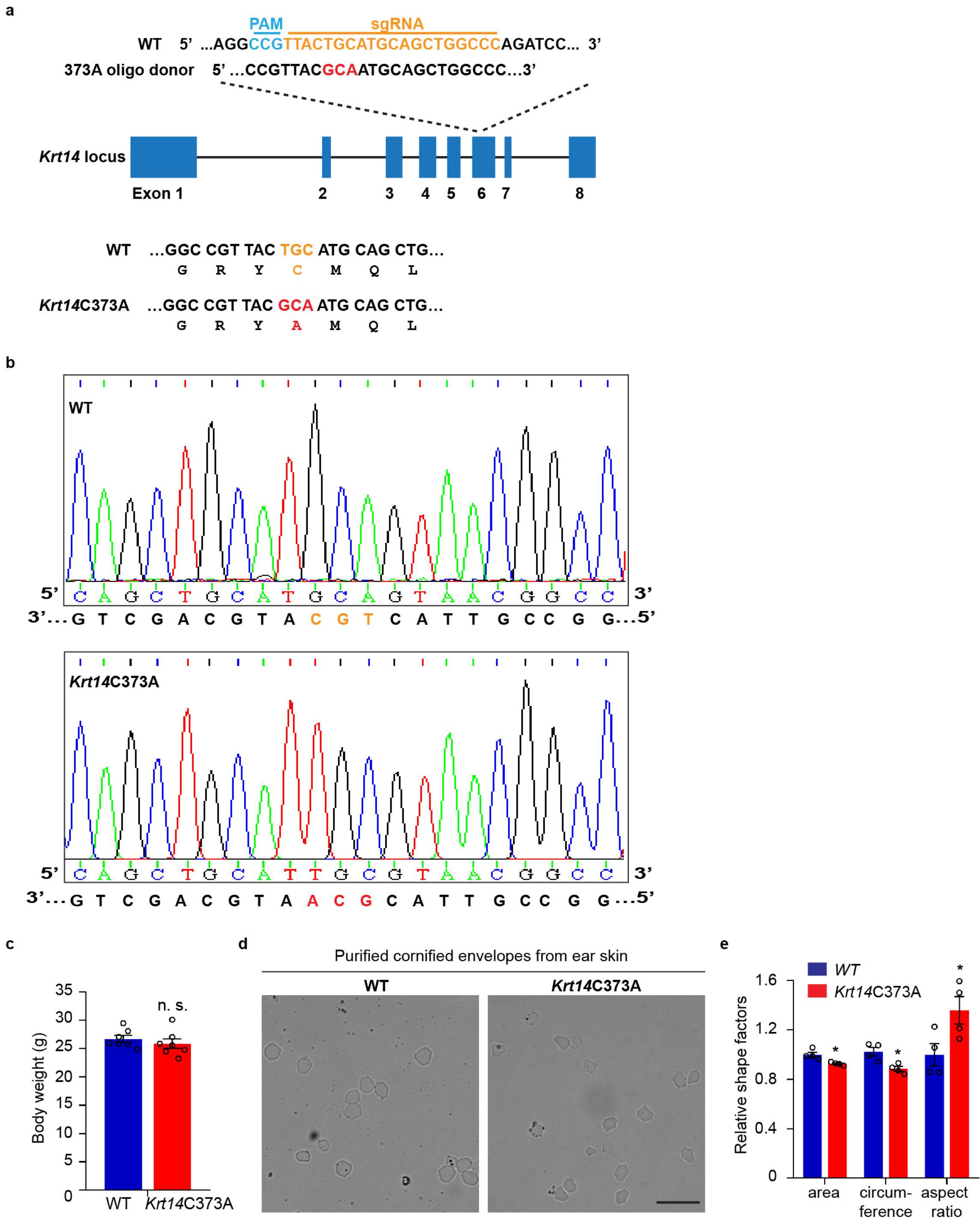

Guo et al. Supplementary Figure 2

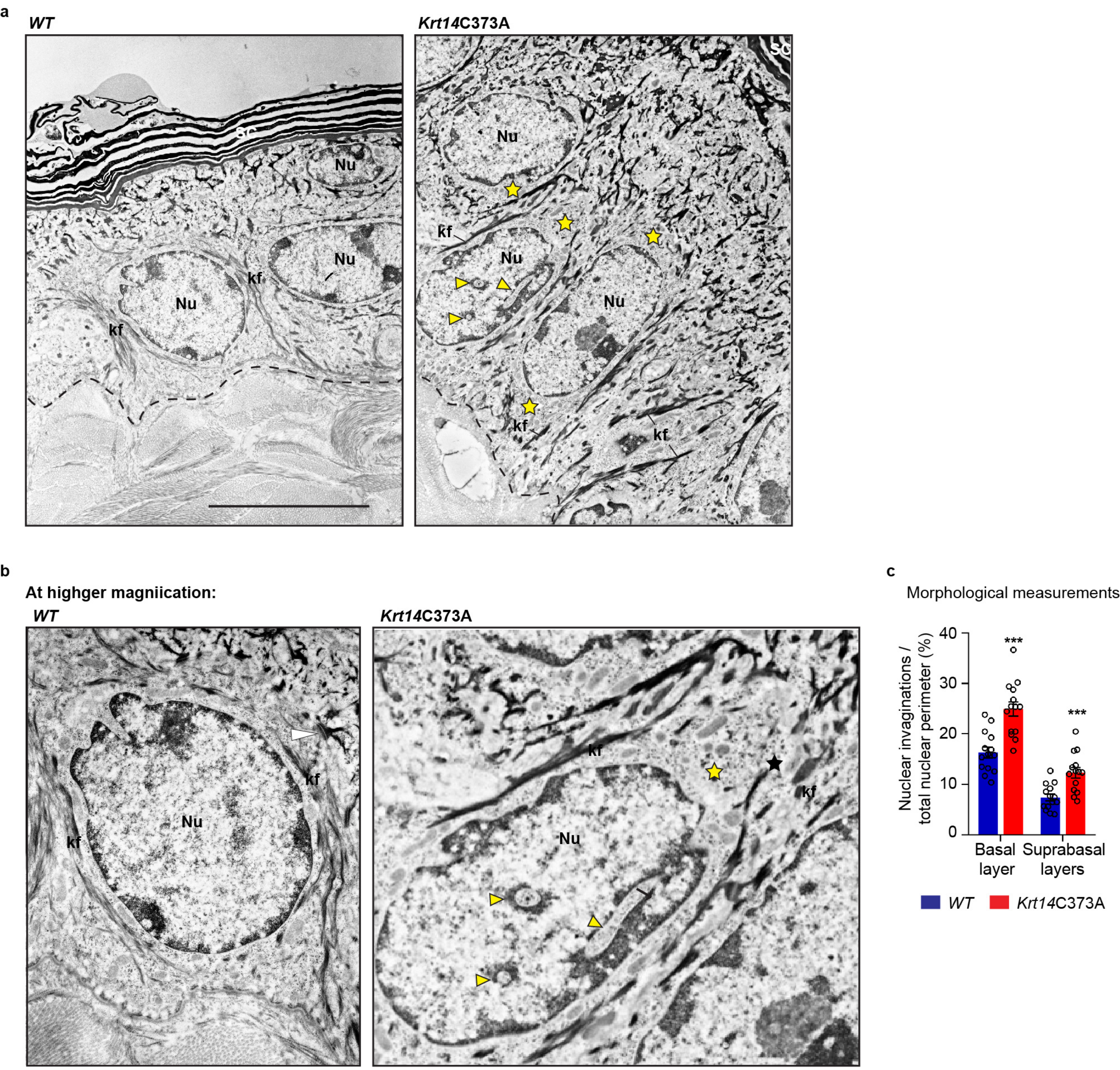

Guo et al., Supplementary Figure 3

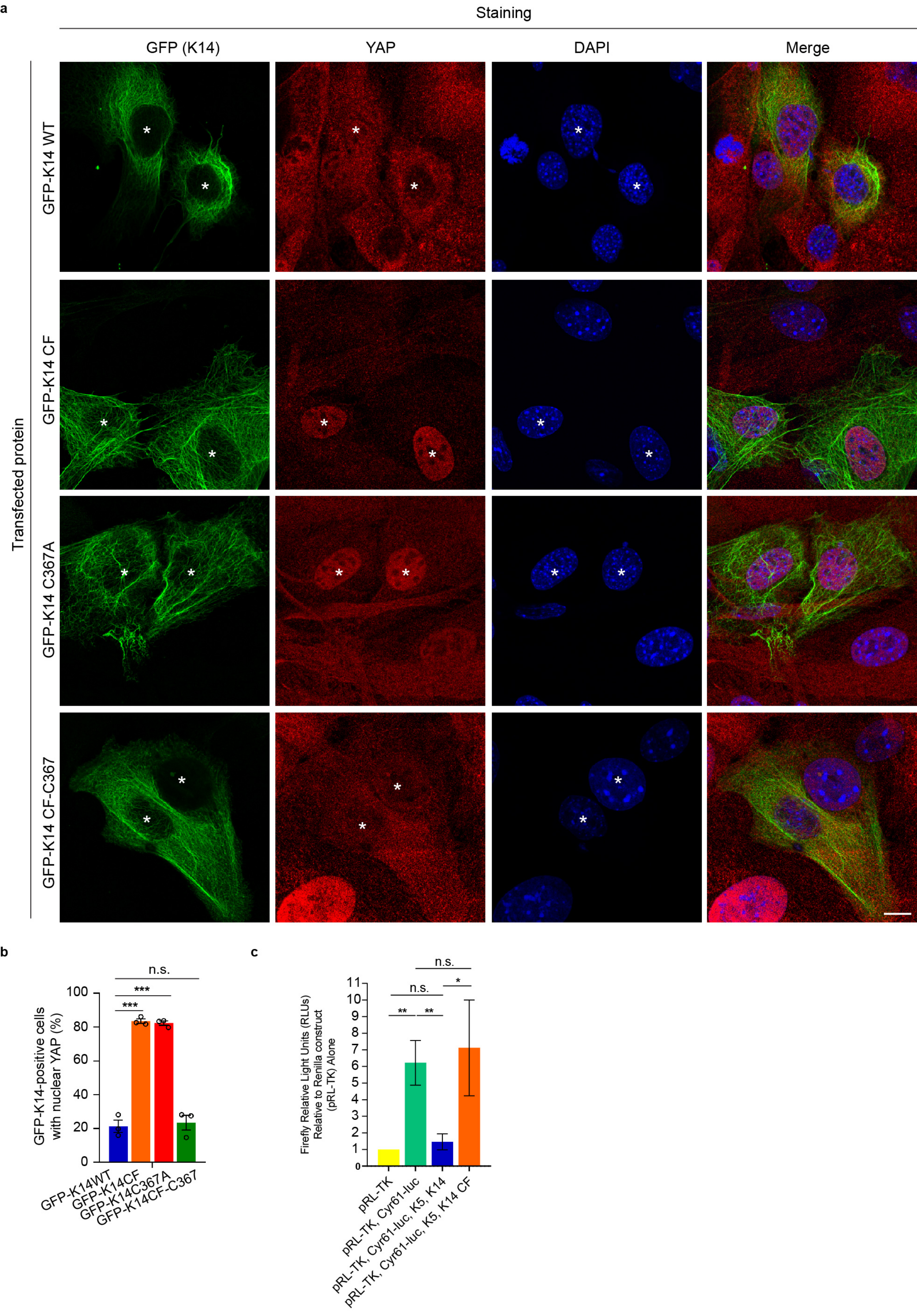

| Gene | Identified Proteins | Accession Number | MW | WT1 | WT2 | WT3 | WT4 | WT5 | Total Counts |
|---|---|---|---|---|---|---|---|---|---|
| Krt5 | keratin, type II cytoskeletal 5 [Mus musculus] | NP_081287.1 | 62 kDa | 360 | 379 | 257 | 313 | 116 | 1425 |
| Krt10 | keratin, type I cytoskeletal 10 [Mus musculus] | NP_034790.1 (+1) | 57 kDa | 181 | 487 | 324 | 307 | 96 | 1395 |
| Krt14 | keratin, type I cytoskeletal 14 [Mus musculus] | NP 058654.1 | 53 kDa | 228 | 287 | 157 | 384 | 244 | 1300 |
| Krt17 | keratin, type I cytoskeletal 17 [Mus musculus] | NP_034793.1 | 48 kDa | 132 | 340 | 94 | 330 | 356 | 1252 |
| Krt6A | keratin, type II cytoskeletal 6A [Mus musculus] | NP_032502.3 | 59 kDa | 307 | 284 | 120 | 325 | 172 | 1208 |
| Krt1 | keratin, type II cytoskeletal 1 [Mus musculus] | NP 032499.2 | 66 kDa | 73 | 409 | 297 | 248 | 77 | 1104 |
| Krt42 | keratin, type I cytoskeletal 42 [Mus musculus] | NP_997648.2 | 50 kDa | 54 | 245 | 137 | 252 | 71 | 759 |
| Krt2 | keratin, type II cytoskeletal 2 epidermal [Mus musculus] | NP_034798.2 (+1) | 71 kDa | 59 | 312 | 145 | 183 | 47 | 746 |
| Krt77 | keratin, type II cytoskeletal 1b [Mus musculus] | NP 032499.2 | 61 kDa | 82 | 246 | 189 | 156 | 0 | 673 |
| Krt16 | PREDICTED: keratin, type I cytoskeletal 16 isoform X1 [Mus musculus] | NP_001300887.1(+1) | 52 kDa | 77 | 154 | 86 | 200 | 76 | 593 |
| Krt19 | keratin, type I cytoskeletal 19 [Mus musculus] | NP_032497.1 | 45 kDa | | | 27 | | 377 | 404 |
| SFN | 14-3-3 protein sigma [Mus musculus] | NP_061224.2 | 28 kDa | 0 | 0 | 254 | 133 | 0 | 387 |
| RPS3 | 40S ribosomal protein S3 (RPS3) [Mus musculus] | NP_058655.3 | 27 kDa | 0 | 232 | 112 | 0 | 0 | 344 |
| Krt15 | keratin, type I cytoskeletal 15 [Mus musculus] | NP_032495.2 | 49 kDa | 0 | 0 | 0 | 194 | 146 | 340 |
| RACK1 | Receptor of activated protein C kinase 1 (RACK 1) [Mus musculus] | NP_032169.1 | 35 kDa | 0 | 273 | 44 | 4 | 0 | 321 |
| DSP | desmoplakin [Mus musculus] | NP_076331.2 (+1) | 333 kDa | 28 | 185 | | 104 | | 317 |
| HspB1 | heat shock protein beta-1 (Hsp27) [Mus musculus] | NP_038588.2 | 23 kDa | 0 | 0 | | 284 | | 284 |
| serpinB6 | serpin B6 isoform a [Mus musculus] | NP_001157589.1 (+7) | 45 kDa | | | 0 | | 258 | 258 |
| ACTB | actin, cytoplasmic 1 [Mus musculus] | NP_031419.1(+6) | 42 kDa | | | 4 | | 253 | 257 |
| Krt75 | keratin, type II cytoskeletal 75 [Mus musculus] | NP_579935.1 | 60 kDa | 0 | 66 | | 186 | | 252 |
| YWHAZ | 14-3-3 protein zeta/delta isoform 1 [Mus musculus] | NP_001240734.1 (+4) | 28 kDa | 0 | 0 | 100 | 149 | 0 | 249 |
| YWHAG | 14-3-3 protein gamma [Mus musculus] | NP_061359.2 | 28 kDa | 0 | 0 | 133 | 96 | 0 | 229 |
| Esd | S-formylglutathione hydrolase isoform 2 [Mus musculus] | NP 001272352.1 (+2) | 31 kDa | 0 | 224 | | 0 | | 224 |
| LMO7 | PREDICTED: LIM domain only protein 7 isoform X12 [Mus musculus] | XP_006519251.1 (+8) | 157 kDa | 212 | 0 | | 0 | | 212 |
| GSTO1 | glutathione S-transferase omega-1 [Mus musculus] | NP_034492.1 | 27 kDa | 0 | 8 | 162 | 23 | 0 | 193 |
| Krt79 | keratin, type II cytoskeletal 79 [Mus musculus] | NP_666175.1 | 58 kDa | 57 | 103 | | 30 | | 190 |
| CAPZB | F-actin-capping protein subunit beta isoform c [Mus musculus] | NP_001258334.1 (+1) | 29 kDa | 0 | 183 | | 0 | | 183 |
| JUP | junction plakoglobin [Mus musculus] | NP_034723.1 (+2) | 82 kDa | 22 | 91 | | 69 | | 182 |
| PRMT1 | protein arginine N-methyltransferase 1 isoform 2 [Mus musculus] | NP_001239405.1 (+3) | 41 kDa | | | 0 | | 179 | 179 |
| HSD17B12 | estradiol 17-beta-dehydrogenase 12 [Mus musculus] | NP_062631.1 | 35 kDa | 0 | 0 | 132 | 44 | 0 | 176 |
| SLC25A3 | phosphate carrier protein, mitochondrial precursor [Mus musculus] | NP_598429.1(+1) | 40 kDa | 0 | 90 | 69 | 0 | 0 | 159 |
| Krt13 | keratin, type I cytoskeletal 13 [Mus musculus] | NP_034792.1 | 48 kDa | | | 154 | | 0 | 154 |
| EIF6 | eukaryotic translation initiation factor 6 [Mus musculus] | NP_034709.1 | 27 kDa | 0 | 0 | | 152 | | 152 |
| DSg1b | desmoglein-1-beta precursor [Mus musculus] | NP_859010.1 (+1) | 114 kDa | 48 | 46 | 21 | 33 | 0 | 148 |
| YWHAB | 14-3-3 protein beta/alpha [Mus musculus] | NP_061223.2 (+1) | 28 kDa | 0 | 0 | 59 | 86 | 0 | 145 |
| YWHAQ | 14-3-3 protein theta [Mus musculus] | NP_035869.1 | 28 kDa | 0 | 0 | 64 | 72 | 0 | 136 |
| SLC25A5 | ADP/ATP translocase 2 [Mus musculus] | NP 031477.1 | 33 kDa | 0 | 0 | 65 | 67 | 0 | 132 |
| RAN | GTP-binding nuclear protein Ran [Mus musculus] | NP_033417.1 (+2) | 24 kDa | 0 | 8 | | 122 | | 130 |
| GAPDH | PREDICTED: glyceraldehyde-3-phosphate dehydrogenase-like isoform | NP_001276655.1 (+3) | 36 kDa | 12 | 91 | | 25 | | 128 |
| CA13 | carbonic anhydrase 13 [Mus musculus] | NP_078771.1 | 30 kDa | 0 | 0 | 20 | 95 | 0 | 115 |
| Krt78 | keratin Kb40 [Mus musculus] | NP_997652.4 (+2) | 112 kDa | 20 | 55 | | 39 | | 114 |
| serpinB2 | plasminogen activator inhibitor 2, macrophage [Mus musculus] | NP_001167641.1 (+2) | 46 kDa | | | 0 | | 112 | 112 |
| RPS27A | ubiquitin-40S ribosomal protein S27a precursor [Mus musculus] | NP_001029037.1 (+8) | 18 kDa | 23 | 18 | 17 | 40 | 13 | 111 |
| YWHAH | 14-3-3 protein eta [Mus musculus] | NP_035868.1 | 28 kDa | 0 | 0 | | 103 | | 103 |
| YWHAE | 14-3-3 protein epsilon [Mus musculus] | NP_033562.3 | 29 kDa | | | 102 | | 0 | 102 |
| PHB | prohibitin [Mus musculus] | NP_032857.1 | 30 kDa | 0 | 0 | 81 | 21 | 0 | 102 |
| RAB25 | ras-related protein Rab-25 [Mus musculus] | NP_058595.2 | 23 kDa | 0 | 0 | | 100 | | 100 |
| RAB5C | ras-related protein Rab-5C [Mus musculus] | NP_077776.2 | 23 kDa | 0 | 0 | | 99 | | 99 |
| ALDOA | fructose-bisphosphate aldolase A isoform 1 precursor [Mus musculus] | NP_001170778.1 (+10) | 45 kDa | 30 | 4 | 0 | 0 | 65 | 99 |
| DHX9 | ATP-dependent RNA helicase A [Mus musculus] | XP 011246220.1 (+1) | 150 kDa | 95 | 0 | | 0 | | 95 |
| GET4 | Golgi to ER traffic protein 4 homolog isoform 1 [Mus musculus] | NP_080545.2 (+1) | 37 kDa | 0 | 92 | | 0 | | 92 |
| TBCB | tubulin-folding cofactor B [Mus musculus] | NP_079824.2 | 27 kDa | 0 | 69 | 21 | 0 | 0 | 90 |
| RAB5A | ras-related protein Rab-5A [Mus musculus] | NP_080163.1 | 24 kDa | 0 | 0 | | 89 | | 89 |
| ARPC2 | actin-related protein 2/3 complex subunit 2 [Mus musculus] | NP_083987.1(+1) | 34 kDa | 0 | 88 | | 0 | | 88 |
| RAB5B | ras-related protein Rab-5B [Mus musculus] | NP_803130.1(+1) | 24 kDa | 0 | 0 | | 85 | | 85 |
| FAM83H | protein FAM83H [Mus musculus] | NP_001161725.1 (+3) | 131 kDa | 83 | 0 | | 0 | | 83 |
| TIM44 | mitochondrial import inner membrane translocase subunit TIM44 [Mus m | NP_035722.2 | 51 kDa | | | 0 | | 81 | 81 |
| RAB21 | ras-related protein Rab-21 [Mus musculus] | NP_077774.1 (+3) | 24 kDa | 0 | 0 | | 79 | | 79 |
| CAPG | macrophage-capping protein [Mus musculus] | NP_001035999.1 (+4) | 39 kDa | | | 0 | | 72 | 72 |
| PKP1 | plakophilin-1 [Mus musculus] | NP_001300630.1 (+1) | 81 kDa | 0 | 47 | | 23 | | 70 |
| ANXA2 | annexin A2 [Mus musculus] | NP_031611.1 (+1) | 39 kDa | 0 | 39 | 11 | 19 | 0 | 69 |
| CYB5R3 | NADH-cytochrome b5 reductase 3 [Mus musculus] | NP_084063.1 (+2) | 34 kDa | 0 | 66 | | 0 | | 66 |
| SLC25A4 | ADP/ATP translocase 1 [Mus musculus] | NP 031476.3 (+1) | 33 kDa | 0 | 0 | | 66 | | 66 |
| serpinB5 | serpin B5 [Mus musculus] | NP_033283.1 | 42 kDa | | | 0 | | 65 | 65 |
| PCBP1 | poly(rC)-binding protein 1 [Mus musculus] | NP 035995.1 | 37 kDa | | | 0 | | 65 | 65 |
| WDR61 | WD repeat-containing protein 61 isoform a [Mus musculus] | NP_001020546.1 (+1) | 34 kDa | 0 | 64 | | 0 | | 64 |
| PHB2 | prohibitin-2 [Mus musculus] | NP_031557.2 | 33 kDa | 0 | 62 | | 0 | | 62 |
| C1QBP | complement component 1 Q subcomponent-binding protein, mitochondri | NP_031599.2 | 31 kDa | | | 62 | | 0 | 62 |
| BUB3 | mitotic checkpoint protein BUB3 [Mus musculus] | NP_001304279.1 (+1) | 37 kDa | | | 0 | | 61 | 61 |
| BLVRA | biliverdin reductase A precursor [Mus musculus] | NP_080954.4 (+3) | 34 kDa | 0 | 55 | | 0 | | 55 |
| HNRNPA3 | heterogeneous nuclear ribonucleoprotein A3 isoform c [Mus musculus] | NP_444493.1 (+3) | 37 kDa | 0 | 29 | 0 | 0 | 26 | 55 |
| ECE-2 | endothelin-converting enzyme 2 isoform d [Mus musculus] | NP_647454.2 | 29 kDa | 0 | 54 | | 0 | | 54 |
| RAB35 | ras-related protein Rab-35 [Mus musculus] | NP_937806.1 | 23 kDa | 0 | 0 | | 53 | | 53 |
| IARS | isoleucine--tRNA ligase, cytoplasmic [Mus musculus] | NP_742012.2 (+1) | 144 kDa | 52 | 0 | | 0 | | 52 |
| Kb14 | type II keratin Kb14 [Mus musculus] | NP_001003670.1 | 60 kDa | 0 | 0 | | 52 | | 52 |
| TGM1 | protein-glutamine gamma-glutamyltransferase K [Mus musculus] | NP 001155186.1 (+4) | 90 kDa | 12 | 31 | | 8 | | 51 |
| MTAP | S-methyl-5'-thioadenosine phosphorylase [Mus musculus] | NP_077753.1 | 31 kDa | 0 | 0 | 35 | 15 | 0 | 50 |

| | | | | | | | | | |
|---|---|---|---|---|---|---|---|---|---|
| HNRNPA1 | heterogeneous nuclear ribonucleoprotein A1 isoform a [Mus musculus] | NP_001034218.1 (+1) | 34 kDa | 0 | 47 | | 0 | | **47** |
| IMPA2 | inositol monophosphatase 2 [Mus musculus] | NP_444491.1 | 32 kDa | 0 | 27 | 20 | 0 | 0 | **47** |
| DPM1 | dolichol-phosphate mannosyltransferase subunit 1 [Mus musculus] | NP_001297013.1 (+1) | 29 kDa | 0 | 0 | | 46 | | **46** |
| MLF2 | myeloid leukemia factor 2 [Mus musculus] | NP_663360.1(+1) | 28 kDa | 0 | 29 | 17 | 0 | 0 | **46** |
| UCHL3 | ubiquitin carboxyl-terminal hydrolase isozyme L3 [Mus musculus] | NP_057932.2 | 26 kDa | 0 | 0 | | 45 | | **45** |
| SBDS | ribosome maturation protein SBDS [Mus musculus] | NP_075737.1 | 29 kDa | 0 | 0 | 26 | 19 | 0 | **45** |
| DNAJB6 | dnaJ homolog subfamily B member 6 isoform a [Mus musculus] | NP_001033029.1 (+4) | 27 kDa | 0 | 0 | | 43 | | **43** |
| SRPRB | signal recognition particle receptor subunit beta [Mus musculus] | NP_033301.1 | 30 kDa | | | 42 | | 0 | **42** |
| MAGT1 | magnesium transporter protein 1 [Mus musculus] | NP_001177338.1(+1) | 42 kDa | 0 | 0 | 18 | 24 | 0 | **42** |
| HSD17B7 | 3-keto-steroid reductase [Mus musculus] | NP_034606.3 (+1) | 35 kDa | 0 | 40 | | 0 | | **40** |
| RAB10 | ras-related protein Rab-10 [Mus musculus] | NP_057885.1 | 23 kDa | 0 | 0 | | 39 | | **39** |
| RAB8A | ras-related protein Rab-8A [Mus musculus] | NP_075615.2 | 24 kDa | 0 | 0 | | 38 | | **38** |
| TPI1 | triosephosphate isomerase [Mus musculus] | NP_033441.2 | 32 kDa | 0 | 0 | | 38 | | **38** |
| ARHGDIB | rho GDP-dissociation inhibitor 2 isoform 1 [Mus musculus] | NP_001288230.1 (+4) | 23 kDa | 0 | 0 | | 37 | | **37** |
| NUDT5 | ADP-sugar pyrophosphatase [Mus musculus] | NP_058614.1 (+4) | 29 kDa | 0 | 37 | | 0 | | **37** |
| RAB6A | ras-related protein Rab-6A isoform 1 [Mus musculus] | NP_001157135.1 (+1) | 24 kDa | 0 | 0 | | 37 | | **37** |
| ANXA1 | annexin A5 [Mus musculus] | NP_033803.1 | 36 kDa | 0 | 36 | | 0 | | **36** |
| RPS4 | 40S ribosomal protein S4, X isoform [Mus musculus] | NP_033120.1(+1) | 30 kDa | 0 | 0 | | 36 | | **36** |
| SRM | spermidine synthase [Mus musculus] | NP 033298.1 | 34 kDa | 0 | 36 | | 0 | | **36** |
| RDH11 | retinol dehydrogenase 11 precursor [Mus musculus] | NP_067532.2 | 35 kDa | | | 36 | | 0 | **36** |
| HAL | histidine ammonia-lyase [Mus musculus] | NP_034531.1 | 72 kDa | 0 | 35 | | 0 | | **35** |
| RRAS2 | ras-related protein R-Ras2 precursor [Mus musculus] | NP_080122.2 | 23 kDa | 0 | 0 | | 35 | | **35** |

#### Supplementary Table 2

**List of antibodies - Guo *et al.***

| Antibody target | Company | Catalogue number | Applications* |
|---|---|---|---|
| K14 | Biolegend | 906001 | IHC, IF |
| K14 | Biolegend | 905301 | WB, IHC, IF, IP, PLA |
| K14 | Abcam | ab7800 | IF and PLA |
| K5 | Biolegend | 905504 | IHC, IF and WB |
| β-actin | Sigma-Aldrich | A5441 | WB |
| K10 | Biolegend | 905401 | IHC |
| Filaggrin | Biolegend | 905801 | IHC and WB |
| Loricrin | Biolegend | 905101 | IHC and WB |
| 14-3-3σ | Santa Cruz Biotechnology Inc | sc-7683 | IHC, WB and IF |
| 14-3-3σ | Sigma-Aldrich | PLA0201 | IF |
| YAP | Santa Cruz Biotechnology Inc | sc-101199 | IHC, IF, WB and PLA |
| Phospho-YAP | Cell Signaling Technology | 4911 | WB |
| YAP | Cell Signaling Technology | 4912 | WB |
| HA | Thermo Fisher Scientific | 26183 | IF and WB |
| HA | Sigma-Aldrich | H6908 | WB |
| α-E-Catenin | Cell Signaling Technology | 3236S | IHC and IF |
| Lamin A/C | Santa Cruz Biotechnology Inc | sc-6215 | IHC and IF |
| Desmoglein 1 | Progen | 61002 | IHC and IF |
| Phospho-MLC2 (Ser19) | Cell Signaling Technology | 3671S | IF |
| Vinculin | Millipore | MAB3574 | IF |
| Alexa Fluor 488 Goat Anti-Mouse IgG (H+L) | Thermo Fisher Scientific | A-11001 | IF |
| Alexa Fluor 488 Goat Anti-Rabbit IgG (H+L) Antibody | Thermo Fisher Scientific | A-11008 | IF |
| Alexa Fluor 546 Goat Anti-Rabbit IgG (H+L) | Thermo FisherScientific | A-11010 | IF |
| Alexa Fluor 555 Donkey Anti-Mouse IgG (H+L) | Thermo Fisher Scientific | A-31570 | IF |
| Alexa Fluor 488 Goat anti chicken IgY (H+L) | Thermo Fisher Scientific | A11039 | IF |

* IF, immunofluoprescence (cells); IHC, immunohistochemistry (tissue sections); IP, immunoprecipitation; PLA, proximity ligation assay; WB, western blotting

**Antibodies used for supplemental data - Guo et al.**

| Antibody target | Company | Catalogue number | Applications |
|---|---|---|---|
| E-cadherin | Cell Signaling Technology | 3195S | IHC |
| Claudin 3 | Thermo Fisher Scientific | 34-1700 | IHC |
| Desmoplakin | K. Green, Northwestern University, Evanston, IL | | IHC |

**Supplementary Table 1 - List of oligonucleotide primers - Guo et al.**

**Primers for analysis of gene expression at the mRNA transcript level**

| Gene | Orientation | Sequence (5'->3') |
|---|---|---|
| *S100A1* | forward | AATGTGTTCCATGCCCATTCG |
| | reverse | ACCAGCACAACATACTCCTTG |
| *S100A2* | forward | GAACAACCTCGATAAGGACAGTG |
| | reverse | CCAGATAGCAGAATCCACCAGA |
| *S100A7A* | forward | AGGAGTTGAAAGCTCTGCTCT |
| | reverse | GCTCTGTGATGTAGTATGGCTG |
| *S100A8* | forward | AAATCACCATGCCCTCTACAAG |
| | reverse | CCCACTTTTATCACCATCGCAA |
| *S100A9* | forward | ATACTCTAGGAAGGAAGGACACC |
| | reverse | TCCATGATGTCATTTATGAGGGC |
| *S100A10* | forward | TGGAAACCATGATGCTTACGTT |
| | reverse | GAAGCCCACTTTGCCATCTC |
| *IL-1a* | forward | CGAAGACTACAGTTCTGCCATT |
| | reverse | GACGTTTCAGAGGTTCTCAGAG |
| *IL-1b* | forward | GAAATGCCACCTTTTGACAGTG |
| | reverse | CTGGATGCTCTCATCAGGACA |
| *HMGB1* | forward | GGCGAGCATCCTGGCTTATC |
| | reverse | GGCTGCTTGTCATCTGCTG |
| *DefB3* | forward | CATCTGCCTCCTTTCCTCAA |
| | reverse | CTTTGCATTTCTCCTGGTGC |
| *TSLP* | forward | ACGGATGGGGCTAACTTACAA |
| | reverse | AGTCCTCGATTTGCTCGAACT |
| *Hspb1* | forward | GGTTGCCCGATGAGTGGTC |
| | reverse | CTGAGCTGTCGGTTGAGCG |
| *Hspa8* | forward | TCTCGGCACCACCTACTCC |
| | reverse | CTACGCCCGATCAGACGTTT |
| *Sprr2d* | forward | GTGGGCACACAGGTGGAG |
| | reverse | GCCGAGACTACTTTGGAGAAC |
| *Sprr3* | forward | CAGGAACCACGGTACGATCT |
| | reverse | TTCCTGGACCATGCTCTACC |
| *MMP9* | forward | CTGGACAGCCAGACACTAAAG |
| | reverse | CTCGCGGCAAGTCTTCAGAG |
| *Ptgs2* | forward | TGC ACT ATG GTT ACA AAA GCT GG |
| | reverse | TCA GGA AGC TCC TTA TTT CCC TT |
| *Cyr61* | forward | CTGCGCTAAACAACTCAACGA |
| | reverse | GCAGATCCCTTTCAGAGCGG |
| *Ctfg* | forward | GGGCCTCTTCTGCGATTTC |
| | reverse | ATCCAGGCAAGTGCATTGGTA |

| | | |
|---|---|---|
| *Zeb1* | forward | TGGCAAGACAACGTGAAAGA |
| | reverse | AACTGGGAAAATGCATCTGG |
| *Snail2* | forward | TGATGCCCAGTCTAGGAAAT |
| | reverse | AGTGAGGGCAAGAGAAAGG |
| *b-actin* | forward | TGGAATCCTGTGGCATCCATGAAAC |
| | reverse | TAAAACGCAGCTCAGTAACAGTCCG |

**Primers for genotyping at the genomic DNA level**

| Gene | Orientation | Sequence (5’-3’) |
|---|---|---|
| *Krt14 WT* | forward | AGACCAAAGGCCGTTACTG |
| | reverse | TTGAGGTGGAGGAGGAGTCT |
| *Krt14* C373A | forward | ACCAAAGGCCGTTACGC |
| | reverse | GAAGCCAAGTCACACCCCTG |
